# Sequential sampling from memory underlies action selection during abstract decision making

**DOI:** 10.1101/2021.04.30.442176

**Authors:** S. Shushruth, Ariel Zylberberg, Michael N Shadlen

## Abstract

The study of perceptual decision making in monkeys has provided insights into the process by which sensory evidence is integrated towards a decision. When monkeys make decisions with the knowledge of the motor actions the decisions bear upon, the process of evidence integration is instantiated by neurons involved in the selection of said actions. It is less clear how monkeys make decisions when unaware of the actions required to communicate their choice—what we refer to as ‘abstract’ decisions. We investigated this by training monkeys to associate the direction of motion of a noisy random-dot display with the color of two targets. Crucially, the targets were displayed at unpredictable locations after the motion stimulus was extinguished. We found that monkeys postponed decision formation until the targets were revealed. Neurons in the parietal association area LIP represented the integration of evidence leading to a choice, but as the stimulus was no longer visible, the samples of evidence must have been retrieved from short-term memory. Our results imply that when decisions are temporally unyoked from the motor actions they bear upon, decision formation is protracted until they can be framed in terms of motor actions.

## Introduction

A decision is a commitment to a proposition or plan of action based on evidence, prior knowledge, priorities and value. Perceptual decision-making refers to the class of decisions in which the dominant source of evidence is derived from sensation and in which the decision is a provisional action or a mental assignment to a category. Viewed from the perspective of information processing, perceptual decision-making establishes a compressed distillation of sensory data into distinct categories. Viewed from the perspective of behavior, it effects an intention, satisfying policy objectives such as obtaining reward. These perspectives are naturally connected because we decide about a perceptual category in order to make a choice. For animals, perceptual decisions typically guide foraging and social choices. For humans, perceptual decisions seem like they are about the perception itself, involving no more than an internal report or change in ideation. The study of decision-making in laboratory animals tends to conflate these depictions, perhaps by necessity.

There is recent interest in characterizing the neural processes that underlie decisions about category membership, independent of intention (e.g., [1–4]), what we will refer to as abstract decisions. Categorical labels introduce flexibility to sensorimotor programs [5,6]. For example, one can assign the labels, “A” and “B” or “rightward” and “leftward”, to consolidate motion perceived to the right or left, irrespective of its precise direction or motion strength. These abstract labels then allow for the implementation of flexible action plans such as “press a red button if you see rightward motion”.

The extent to which nonhuman primates can assign abstract labels to sensory percepts and exploit them to be flexible in their actions is unclear. The process of abstraction, by definition, unyokes the sensory evaluation processes from the process of acting on the sensory information. However, multiple lines of research in macaques suggest that the process of sensory evaluation is intimately coupled to the actions that can result from the evaluative process [7–9]. This framework, wherein cognitive processes are embodied in terms of the motor actions they afford, is supported by the patterns of neural activity found in association and premotor cortices of monkeys [7,10,11]. Yet monkeys can be trained to decide on properties of sensory stimuli even when unaware of the exact motor action that will be required of them to report their decision [1,2,4,12–15]. In these studies, monkeys were required to commit to a category assignment without committing to an action. The studies show that monkeys are capable of performing such tasks, but it is unclear how the abstract representation is established and how it is ultimately translated to the response. That is what we set out to clarify.

We trained two monkeys to decide on the net direction of stochastic random dot motion and associate two possible directions with two colors. The monkeys reported the direction of motion by making an eye movement to the target of the associated color, but these targets were revealed at unpredictable locations after the motion stimulus had been extinguished. To perform the task well, monkeys needed to integrate motion information in the stimulus over time to make an abstract decision about the direction of motion. This imposition allowed us to investigate how an abstract perceptual decision is formed when the actions associated with the decision are yet to be specified. Further, since the decision making phase is unyoked from the motor planning phase, the task also permits investigation of the conversion of an abstract decision to an action.

Surprisingly, we found that evidence evaluation and action selection—the two aspects of abstract decision making that our task was supposed to unyoke—were, in fact, intimately coupled. The behavior of the monkeys showed that they based their decision on motion evidence integrated over time. However, this integration transpired during the action selection epoch instead of the epoch when the evidence was presented. Further, activity of neurons in the sensorimotor association area LIP represented decision formation during the action selection epoch. Our results suggest that monkeys form abstract perceptual decisions by evaluating sensory information from iconic short term memory (e.g., [16]) for action selection.

## Results

We trained two monkeys to decide whether the net direction of a random dot motion (RDM) stimulus was to the right or left. The monkeys reported their decision by making an eye movement to a blue or yellow target based on the association they had learned between the direction of motion and target colors (Fig. 1). The two targets appeared after a short delay (200–333 ms) following the termination of the motion stimulus, and the locations of the two targets were randomized across trials. Thus all the evidence bearing on the decision was supplied before the monkeys were instructed about the motor act that would be required to report the decision. Unlike previous studies [12,14], both monkeys were naive to the RDM stimulus when they began training on the task. Since one of our goals was to investigate how decisions are converted to motor actions, the monkeys were allowed to report the decision as soon as the targets were presented (*go*-task). Monkey-SM was also trained on a variant of the task in which an additional waiting time was imposed after the appearance of the targets (*wait*-task).

**Fig 1.**
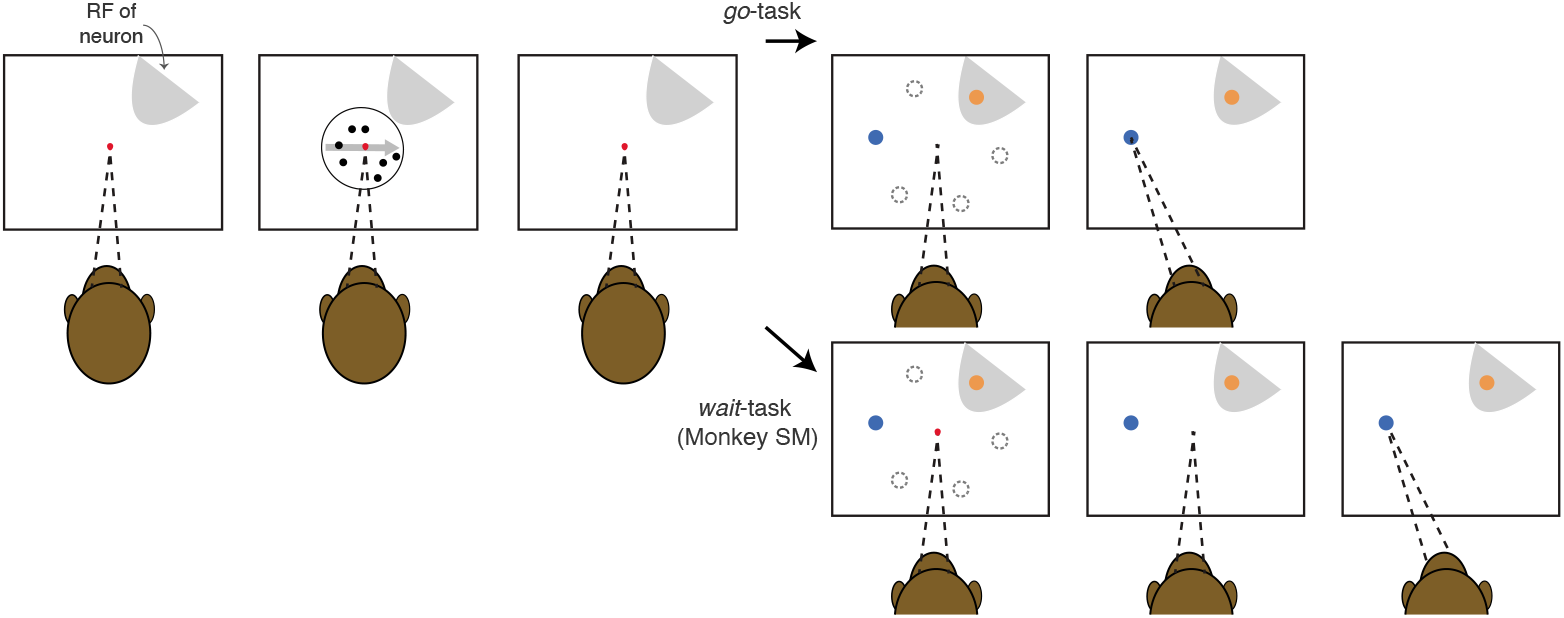
Behavioral task. The monkey fixates at an instructed location (red dot). After a delay, a random dot motion stimulus appears around the fixation point. The stimulus terminates after a variable duration (350–800 ms). After another short delay (200-333 ms), a blue and a yellow target appear at unpredictable peripheral locations. In the *go* version of the task (top panel), the fixation is extinguished at the time the targets appear and the monkey can report the decision by choosing one of the colored targets. In the *wait* version of the task (bottom panel), the monkey must wait until the fixation point is extinguished before choosing a target. During recording sessions, the target locations on each trial are pseudorandomly chosen from a restricted set of locations based on the receptive field of the neuron being recorded. The unchosen locations are illustrated by dashed gray circles (not shown to the monkey). During training sessions, the target locations were less constrained.

The abstract decision-making task proved to be challenging for the monkeys to learn (Supp. Fig. 2-1). Monkey-AN required 28 sessions to acquire the motion-color association, and failed to improve beyond competency at the highest motion strengths for the next ~40 sessions (~50,000 trials). Only then did the monkey begin to exhibit gradual improvement, quantified by a reduction in psychophysical threshold—the motion strength required to support accuracy greater than 75% correct (Eq. 1). Monkey-SM learned the motion-color association quickly but made little progress over months of training. After 127 sessions (983 trials per session on average), the thresholds still hovered around 25% coherence. This monkey was then trained on the wait variant of the task for an additional 58 sessions (740 trials per session) until the thresholds decreased and stabilized at ~11% coherence.

By the final training session, both monkeys performed the task above chance for all non-zero motion coherences (Fig. 2A,B), although they made many errors on the easiest motion strength. Such asymptotic performance is commonly interpreted as a sign that the decision-maker lapses (e.g., guesses) on a fraction of all trials. The lapse rates were 9% and 11% of trials for monkeys AN and SM, respectively. Monkeys performing the same direction discrimination task with a direct mapping between motion direction and actions typically exhibit lapse rates under 2%, further attesting to the challenging nature of the present task, even after extensive training. Nonetheless, for the vast majority of trials, both monkeys used evidence from the RDM to choose the appropriate color. This was confirmed using psychophysical reverse correlation, which identifies the times that random fluctuations of information in the random dots stimulus influence the decision. The analysis reveals that monkeys AN and SM based their decisions on information acquired over 357 ms and 261 ms, respectively (Fig. 2C,D). Importantly, the influence ceases at least 350 ms before the color targets appear.

**Fig 2.**
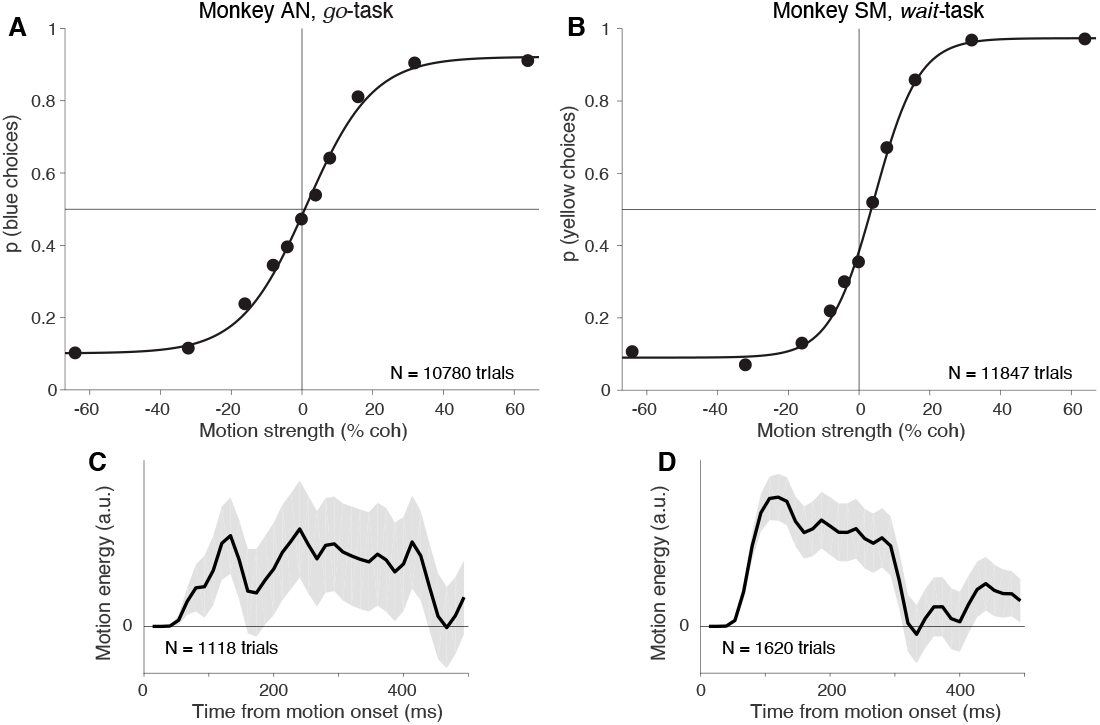
Color choices are governed by the strength and direction of motion. **A**, Effect of motion strength on decisions for monkey-AN on the *go*-task. Monkey-AN was trained to associate blue with rightward and yellow with leftward. The proportion of blue (rightward) choices are plotted as a function of signed motion strength (rightward motion is positive signed). Curves are logistic regression fits to the data. Error bars are s.e. **B**, Effect of motion strength on decisions for monkey-SM on the *wait*-task. Monkey-SM was trained to associate yellow with rightward and blue with leftward. The proportion of yellow (rightward) choices are plotted as a function of signed motion strength. Otherwise, same conventions as in A. **C–D**, The influence of fluctuations in motion information on choices plotted as a function of time from motion onset. Curves represents the mean motion energy in support of the direction chosen by the monkey on 0% coherence trials (shading, ±1 s.e.m.).

### Action selection during abstract decision making is a deliberative process

The behavioral task is structured to separate the decision-making epoch from the action-selection epoch. The natural expectation is that the monkey makes a decision about the direction of motion in the random dot display while it is visible and is thus prepared to make an eye movement to the blue or yellow target once they are displayed (Fig. 3, Strategy 1). If so, the action-selection epoch would involve a simple translation of the decision into an eye movement. This should take as little as 200 ms—the amount of time required to search for a colored target accompanied by a single, highly discriminable distractor [17–19]. Unexpectedly, we found that both monkeys required a prolonged action-selection epoch to integrate motion evidence towards a decision. Given that only the two colored targets were visible during this epoch, we hypothesized that monkeys could be using this time to sample information stored in memory to render their decisions (Fig. 3, Strategy 2). Behavioral results from monkeys AN and SM provided distinct but complementary insights into the deliberative process. We next proceed to describe them separately, followed by the data from neural recordings which were strikingly similar.

**Fig 3.**
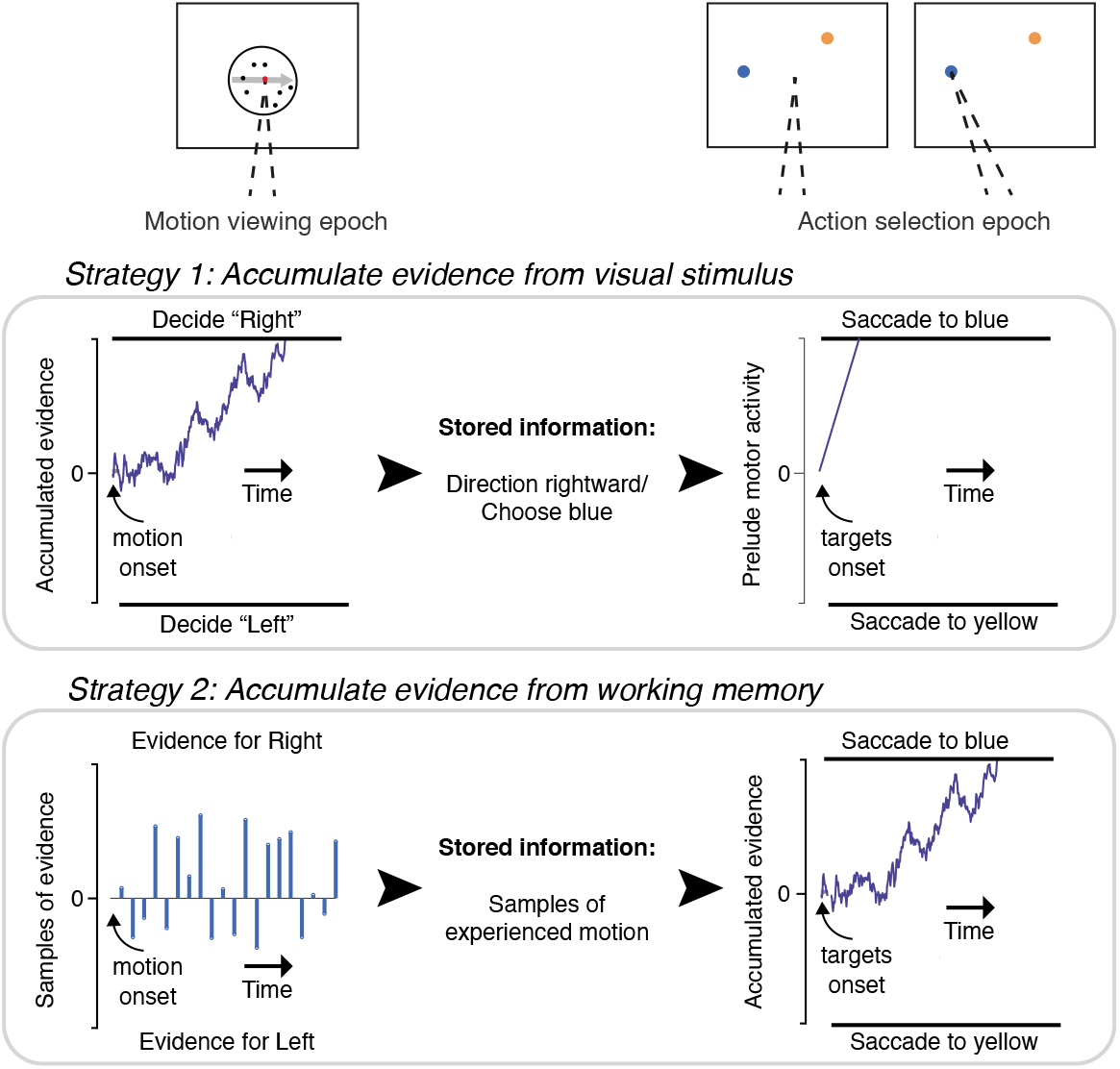
Putative strategies. Schematic of strategies that monkeys could adopt to solve the task. *Strategy 1*: During motion viewing, evidence for motion direction is accumulated to decide if the motion is to the right or left. The result of the decision about direction and/or its color association is stored. When the colored targets are presented, the previously made decision guides an immediate saccade to the target with the chosen color. The saccadic latency might vary by 10–20 ms as a function of confidence in the decision. *Strategy 2*: During motion viewing, the experienced evidence is stored in short term memory. When the targets are shown, the stored evidence is evaluated during action-selection to decide which of the two colored targets to choose. The drawing gives the impression of many samples, but the samples themselves might represent several tens of ms of motion information (as in [20]).

### Monkey-AN: Prolongation of go-RTs

Monkey-AN showed a natural inclination to deliberate after the appearance of the targets. Fig. 4 displays the proportion of blue choices (*bottom*) and their associated response times (go-RT, *top*), as function of motion strength and direction. The response times, measured from onset of the colored choice targets (at least 300 ms after the motion stimulus was extinguished), were 2–4 times slower than expected if the decision had been made during the motion viewing epoch (i.e., Strategy 1). The averages range from 440 ms for the easiest condition to 771 ms for the most difficult, and they exhibit a clear dependency on the strength and direction of motion. The range of go-RTs between easiest to most difficult blue choices and the range between easiest to most difficult yellow choices is an order of magnitude longer than the range associated with changes in reward expectation or confidence (typically < 20*ms*; e.g. [12]). The dependency of go-RT on motion strength resembles the pattern of response times—relative to onset of the RDM—seen in earlier studies, where monkeys were free to indicate their saccadic choices to targets that were already present during motion viewing (e.g., [21]).

**Fig 4.**
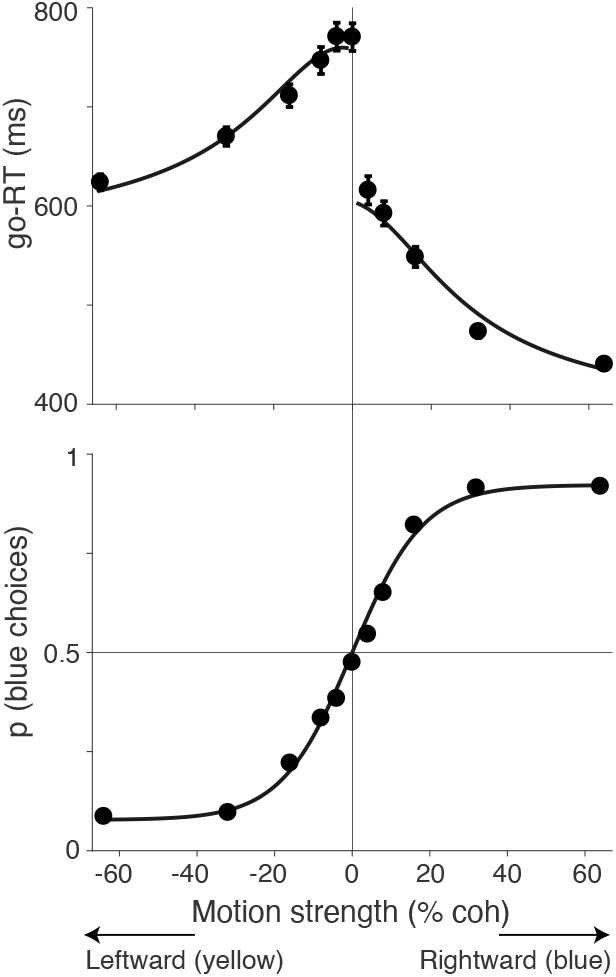
Deliberation during action selection in monkey-AN. *Top,* Go-RTs of monkey-AN plotted as a function of signed motion coherence. Curves are fits to a bounded drift-diffusion model. The model is also constrained by the choice proportions. *Bottom,* Same data as in Fig. 2A. Curve is the fit of the bounded diffusion model, which accounts for both the choice proportions and the go-RTs.

We therefore considered the possibility that the pattern of go-RTs might result from sequential sampling of evidence experienced earlier in the trial, that is, from memory (Fig. 3, Strategy 2). To evaluate this, we appropriated a bounded evidence-accumulation model (drift-diffusion) that is known to reconcile the choice proportions with the response times of subjects when they are allowed to answer whenever ready. In such *free response* tasks, the decision-maker knows how to answer while viewing the motion and simply stops the trial by pushing a button or making an eye movement to one of two visible targets. We wondered if the same type of model could reconcile the choices and go-RTs of monkey-AN.

The curves in Fig. 4 are fits of a bounded drift-diffusion to the proportion of blue choices and the mean go-RTs. The simplest version of this model assumes the drift rate is proportional to signed motion coherence and the terminating bounds do not change as a function of time [22]. Any bias is accommodated by an offset to the drift rate [23]. The mean go-RT for each signed motions strength is predicted by the expectation of the bound termination times plus a constant non-decision time, which captures contributions to the response time that do not depend on the motion strength and bias. We used separate terms (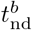 and 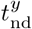) to describe the faster blue and slower yellow choices. The model to this point uses only five degrees of freedom to explain the choice proportions and mean go-RT across 11 motion strengths (Eq. 3 and Eq. 4).

We incorporated one additional feature to accommodate the failure of monkey-AN to achieve perfect performance on the easiest conditions (±64% coh). Such errors are typically attributed to lapses in which the subject ignores the evidence and guesses. However we noticed that the go-RTs associated with errors on the strong leftward condition (blue choices), had the slow go-RTs associated with correct leftward (yellow) choices. Similarly, the errors associated with the strong rightward condition (yellow choices) had the fast go-RTs associated with correct rightward choices (Supp. Fig. 4-1). This indicates that the lapses were not guesses but an error in the association between direction and color. We accommodated this feature in the model, assuming that this type of error occurred on a small fraction of trials, independently of motion strength (see Methods). The model captures the coherence dependence of the go-RTs on correct choices (*R^2^* = 0.99) while also accounting for the accuracy of the monkey’s choices (Fig. 4). The fidelity of the fits supports the hypothesis that the prolonged go-RTs reflect a bounded accumulation of noisy evidence leading to the rendering of the decision. As this sampling began at least 300 ms after the motion stimulus was extinguished, these samples must be derived from memory. In Appendix S1, we show that the wrong association between direction and color provides a better account of the data than an alternative model in which the errors are due to low sensitivity to motion information.

Because drift-diffusion models with time-independent (i.e., ‘flat’) bounds cannot explain the difference in response times between correct and error choices at a given motion strength, we considered a more elaborate version of the model to explain the go-RTs on errors. The model incorporates decision-termination bounds that can change with elapsed time (Eq. 11). We fit the extended model to the choice and go-RT data, including the response times on errors. The best fitting model (Supp. Fig. 4-1) yields an expectation of the integration time for each motion strength. For 0% coherence, the expectation is 243 ms, which is consistent with the psychophysical reverse-correlation analysis, above (Fig. 2C). Note that the reverse correlation analysis also shows that the monkey uses the earliest epochs of motion evidence to inform its decision. Taken together, the analyses of go-RT suggest that monkey-AN stores at least 300 ms of information about the motion in some form. The estimate is longer than the expectations, because the duration of stimulus information needed for decision termination is not known before the accumulation process transpires.

### Monkey-SM: Improved integration after enforced wait

Unlike monkey-AN, monkey-SM did not show a tendency to deliberate after target onset in the go-version of the task. But while monkey-SM learned the direction-color association and performed better than monkey-AN at the strongest motion conditions, it failed to achieve proficiency on the more difficult conditions (Supp. Figure 2-1). Even after extensive training, sensitivity plateaued at an unacceptable level (Fig. 5A, green), and psychophysical reverse correlation revealed only a weak, transient impact of motion information on choice (Fig. 5D). We therefore suspected that this monkey based its decisions on a brief sample of information from the first, last or random glimpse of the display (e.g., see [24]). We confirmed this using a variant of the *go* task in which the strength of motion was modulated as a function of time within a trial (Supp. Fig. 3-2). The coherence started at 0% and either stepped or changed gradually to a large positive or negative value. The time of the step or the rate of change varied across trials. The monkey’s performance deteriorated to chance when the strong motion was concentrated at the end of the trial (Supp. Fig. 5-1). We deduced that the monkey based its decisions on motion information sampled over a short time window at the beginning of the trial. Not surprisingly, the go-RTs exhibited no sign of deliberation. They were nearly as fast as a simple saccadic reaction time to a single target (192 ± 0.4 ms) and showed no influence of the previously experienced motion strength.

**Fig 5.**
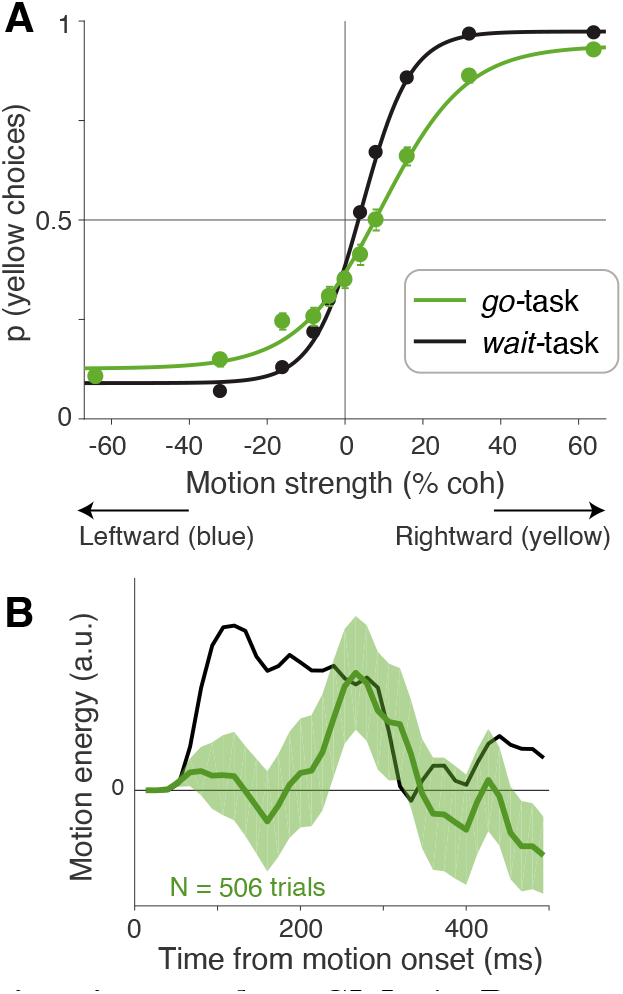
Deliberation during action selection in monkey-SM. **A**, Proportion of rightward (yellow) choices as a function of motion strength for monkey-SM from the last four training sessions on the *go*-task (green). The same data as Fig. 2B is shown for comparison (black). Lines are logistic regression fits to the data. **C**, Influence of motion fluctuations on choice in the last four training sessions of the *go*-task for monkey-SM (green curve). Same conventions as Fig. 2D. Data from the *wait*-task (black curve, same as in Fig. 2D) is shown for comparison.

Based on our experience with monkey-AN, we wondered if monkey-SM failed to integrate after the color-choice targets appeared. We therefore introduced a wait time after the onset of the targets. This simple modification led to a twofold improvement in sensitivity (Fig. 5B, green vs. black traces; also see Supp. Fig. 2-1). This degree of improvement would require at least a fourfold increase in the number of independent samples of evidence the monkey used to form its decision. Indeed, psychophysical reverse correlation revealed a longer time window over which motion information influenced decisions: from 40 ms, before the introduction of the enforced wait, to 261 ms, after ~40 sessions of training (Fig. 5B), Thus, the imposition of a wait after the onset of the targets encouraged monkey-SM to use more information to inform its decision—information that was acquired earlier, in the motion viewing epoch.

The behavioral data from both monkeys therefore provides complementary evidence that deliberation during the action-selection epoch is necessary for integrating previously observed motion information. During motion viewing both monkeys must store some representation of the motion in short term memory. The go-RTs from monkey-AN indicate that the stored information is sampled sequentially in the action selection period. Owing to the enforced wait, we lack meaningful go-RTs for this monkey. However, as we next show, the neural recordings demonstrate that monkey-SM also samples sequentially from memory during the action-selection epoch.

### LIP neurons represent the accumulation of evidence from memory

We recorded from single units with spatially selective persistent activity in area LIP [25, 26]. Such neurons are known to represent an evolving decision variable—the accumulated evidence for and against a motion direction—when one of the choice targets is in the neural response field (RF) [21,27]. The present study differs from previous reports in two critical aspects: (*i*) the choice targets were not visible during motion viewing, and (*ii*) the locations of the choice targets were unpredictable. Under these conditions, the neural responses accompanying motion-viewing were only weakly modulated by motion strength in monkey-AN (Fig. 6A) and unmodulated in monkey-SM (Fig. 6D).

**Fig 6.**
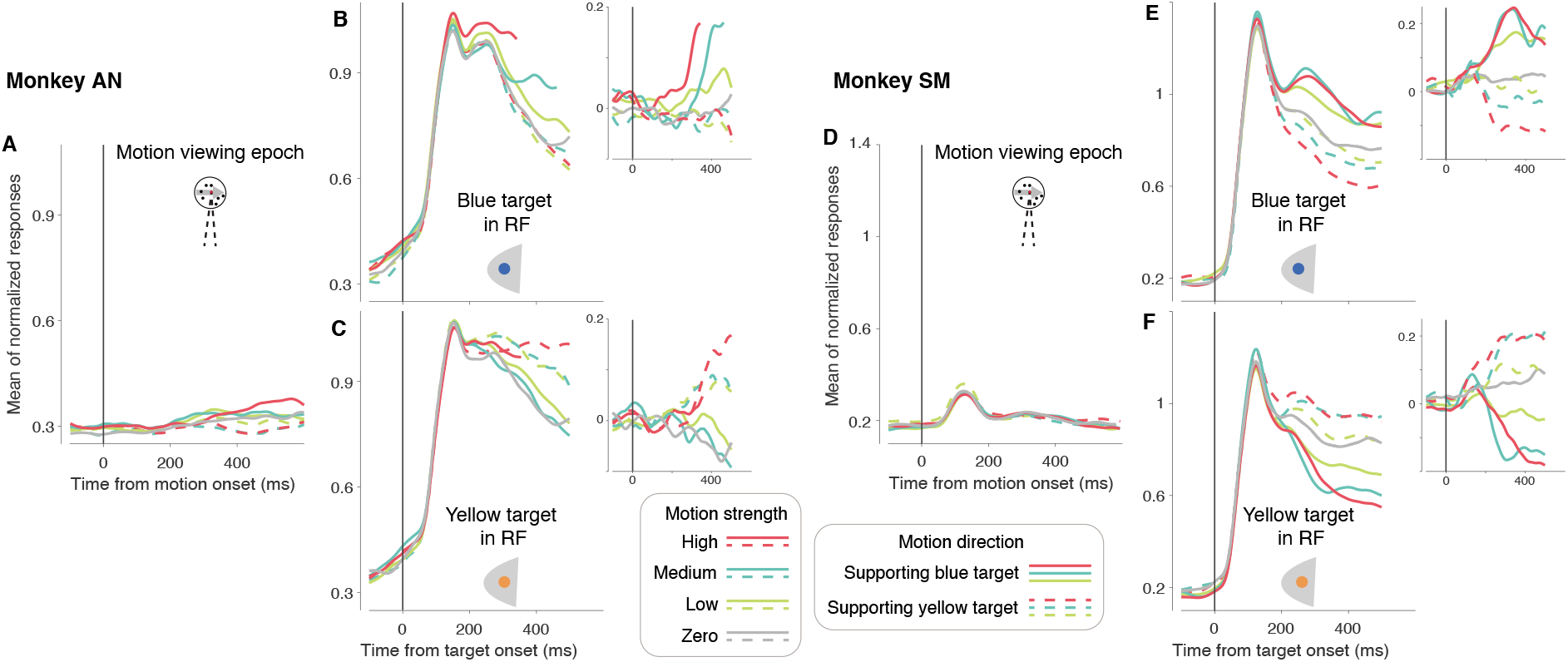
LIP activity during motion viewing and target selection. The graphs show average normalized responses as a function of time aligned to motion onset or target onset. Data from the two monkeys are shown separately (*left,* AN, 29 neurons; *right,* SM, 31 neurons). **A,D** Responses aligned to motion onset. All trials are included. **B,C,E,F** Data from trials in which the blue (B,E) or yellow (C,F) target was in the neuronal response field. The responses are aligned to the onset of the target. Insets show residual responses after removal of the large visual response to the target. They isolate the component of the response that is controlled by the strength and direction of motion. In all panels, coherences are grouped as High (±64% and ±32%), Medium (±16%), Low (±8% and ±4%) and 0%. Grouping of the direction of motion is based on the preferred color-motion association for each neuron. This was consistent with the association the monkey had learned between motion direction and target color, except for six neurons in monkey-SM for which the association was reversed (see Methods). The responses aligned to saccade are shown in Supp. Fig. 6-1.

The action selection epoch begins with the appearance of the color-choice targets. When a target was in the neural RF, it elicited a strong visual response beginning ~50 ms after onset (Fig. 6B,C,E,F), consistent with previous reports [28]. The subsequent evolution of the response reflected both the strength and direction of the RDM stimulus that had been presented in the previous epoch. To better visualize the relationship between the neuronal response and the previously presented RDM stimulus, we removed the visual response (see Methods). The residual responses (Fig. 6B,C,E,F, *insets*) are effectively detrended with respect to any influences that are unaffected by the strength and direction of motion. The residual responses exhibit a clear dependency on the strength and direction of the RDM.

To quantify the rate of change of residual responses (*buildup rate*), we identified the time at which the raw responses first diverge. For each neuron, we then computed the buildup rate for each coherence as the slope of a line fit to the average of the residual firing rates. Each point displayed in Fig. 7 is the mean buildup rate across neurons. These buildup rates exhibited a linear dependence on motion strength. For monkey-SM, the linear dependence was statistically significant in all four conditions (i.e., for all combinations of direction of motion and color of target in the RF; see Table 1). For monkey-AN, the linear dependence was statistically significant in three of the four combinations of motion and direction (*p* < 0.05, Table 1). The magnitude of the dependencies are comparable to those obtained under simpler task, when the motion is viewed in the presence of saccadic choice targets (e.g., see Figure 3G in [29]).

**Fig 7.**
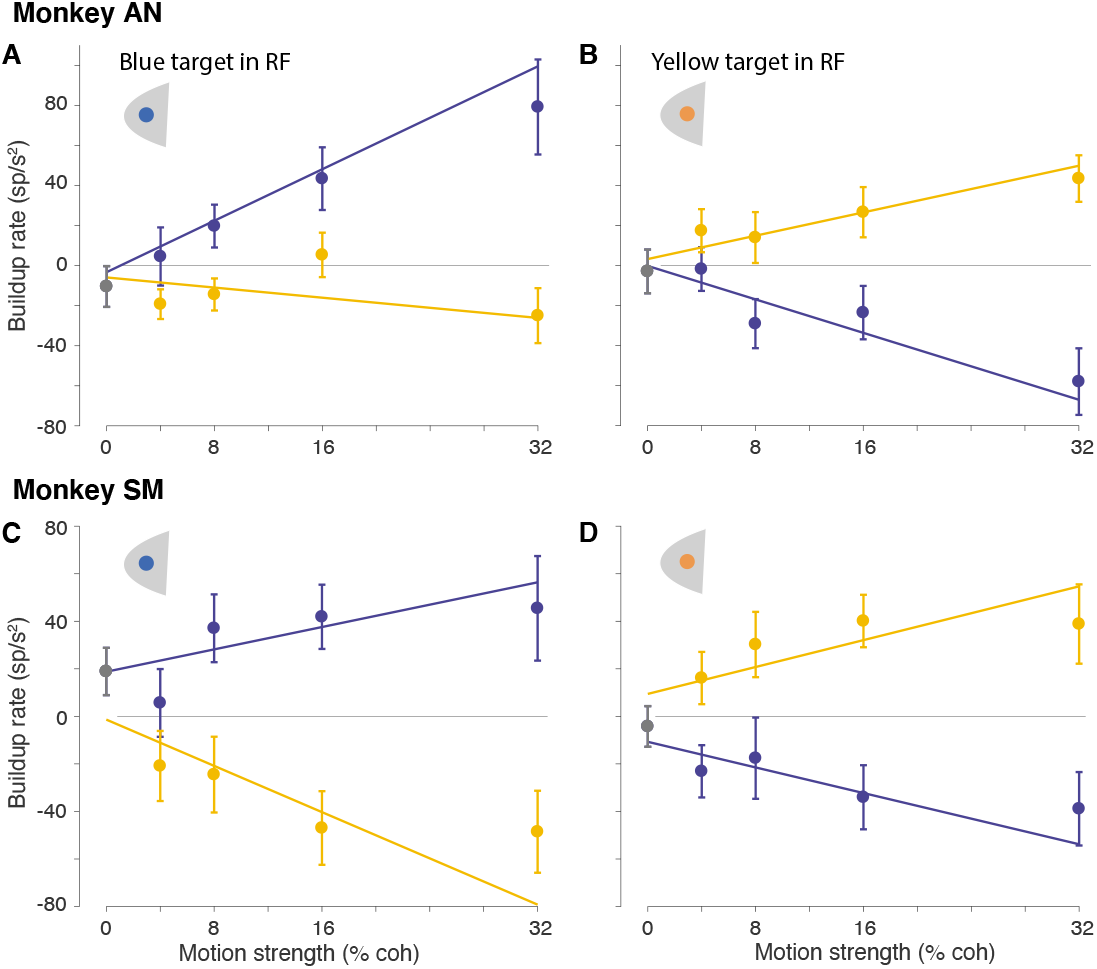
Buildup of neural activity depends on the strength and direction of motion. Buildup rates were estimated for each neuron, using trials with the same motion strength, direction, and color-target in the response field (*top*, monkey-AN; *bottom*, monkey-SM). Symbols are averages across neurons (error bars are s.e.m.) The lines in the graph are weighted least square fits to the average buildup rates, grouped by motion direction. The 0% coherence point (gray) is included in both weighted regressions in each panel.

**Table 1.**
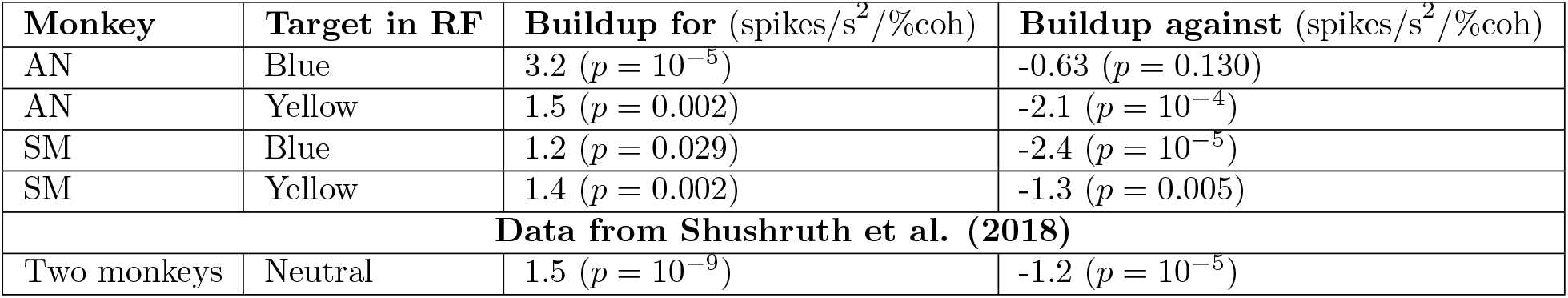
Dependence of buildup rates on motion strength.

Thus for both monkeys, the neural responses during action selection exhibit the hallmark of a decision variable, which must be informed by evidence acquired earlier. This is consistent with the pattern of go-RTs in monkey-AN, which also support sequential sampling of evidence during the action-selection epoch. Analyses of the time dependent changes in response variance and autocorrelation lend additional support for sampling of noisy evidence from memory in both monkeys.

The coherence dependent ramping evident in trial-averaged response residuals could reflect the accumulation of noisy samples of evidence on single trials. The trial-averages suppress the noise, leaving mainly the deterministic component of the accumulation—a ramp with slope equal to the statistical expectation of the momentary evidence for yellow or blue. On single trials, theoretically, the decision variable also includes an accumulation of noise. This is the diffusion component of drift-diffusion, which is thought to explain stochastic choice and variable decision times. Although suppressed in the trial averages, the diffusion component can be detected in the evolution of the variance and autocorrelation of the neural firing rates [29–31]. The procedure utilizes spike counts from single trials, which provide a noisy estimate of the rate over a short counting window. The counts are thus conceived as the result of a doubly stochastic process: a rate that represents a diffusion (or random walk) process, which differs from trial to trial, and the stochastic point process that renders spike-counts from the rate. The strategy is to remove the latter component of the total variance to reveal the variance of the conditional expectation of that count. Hence we refer to variance of the rate, at the time of a counting window, as the variance of the conditional expectation (VarCE) of the count. We adapted this procedure to the current data set in order to ascertain whether the residual firing rates on single trials incorporate the accumulation of independent samples of noise (see Methods, Eq. 12-Eq. 16).

We divided the period following target onset into 60 ms bins and computed the VarCE across trials for each bin. We identified the epoch of putative accumulation to coincide with the time of the buildup. The VarCE underwent a linear increase as a function of time over most of this epoch (Fig. 8A,F. This is the pattern expected for partial sums (i.e., accumulation up to time, *t*) of independent samples of noise. The autocorrelation of the responses (CorCE) also showed signatures of a diffusion process: a decrease in autocorrelation as a function of the time separation between the bins (i.e., lag) and an increase in autocorrelation between adjacent bins as a function of time (Fig. 8C,E and H,J). The estimated autocorrelation pattern for both monkeys hewed closely to the theoretical predictions (*R^2^* = 0.84, monkey-AN, Fig. 8D; *R^2^* = 0.89, monkey-SM, Fig. 8I). Such conformance lends further support for the conclusion that evidence integration in both monkeys occurs in the action-selection epoch. Indeed, the same analyses applied to the neural responses in the motion-viewing epoch fail to conform to the theoretical predictions of diffusion (i.e., integration of noisy evidence) (*R^2^* = 0.45, monkey-AN; *R2* = 0.26, monkey-SM, Supp. Fig. 8-1). This was already obvious from the response averages in Fig. 6A,D. The demonstration of a dynamic process of evidence accumulation rules out other potential strategies that might explain the neural responses. In particular, we considered the possibility that the monkeys form their decisions during motion viewing (using neurons outside of area LIP), and the LIP activity in the action-selection epoch is simply a reflection of confidence in the decision. If so, the neural activity in the action-selection epoch should not represent the accumulation of noisy evidence.

**Fig 8.**
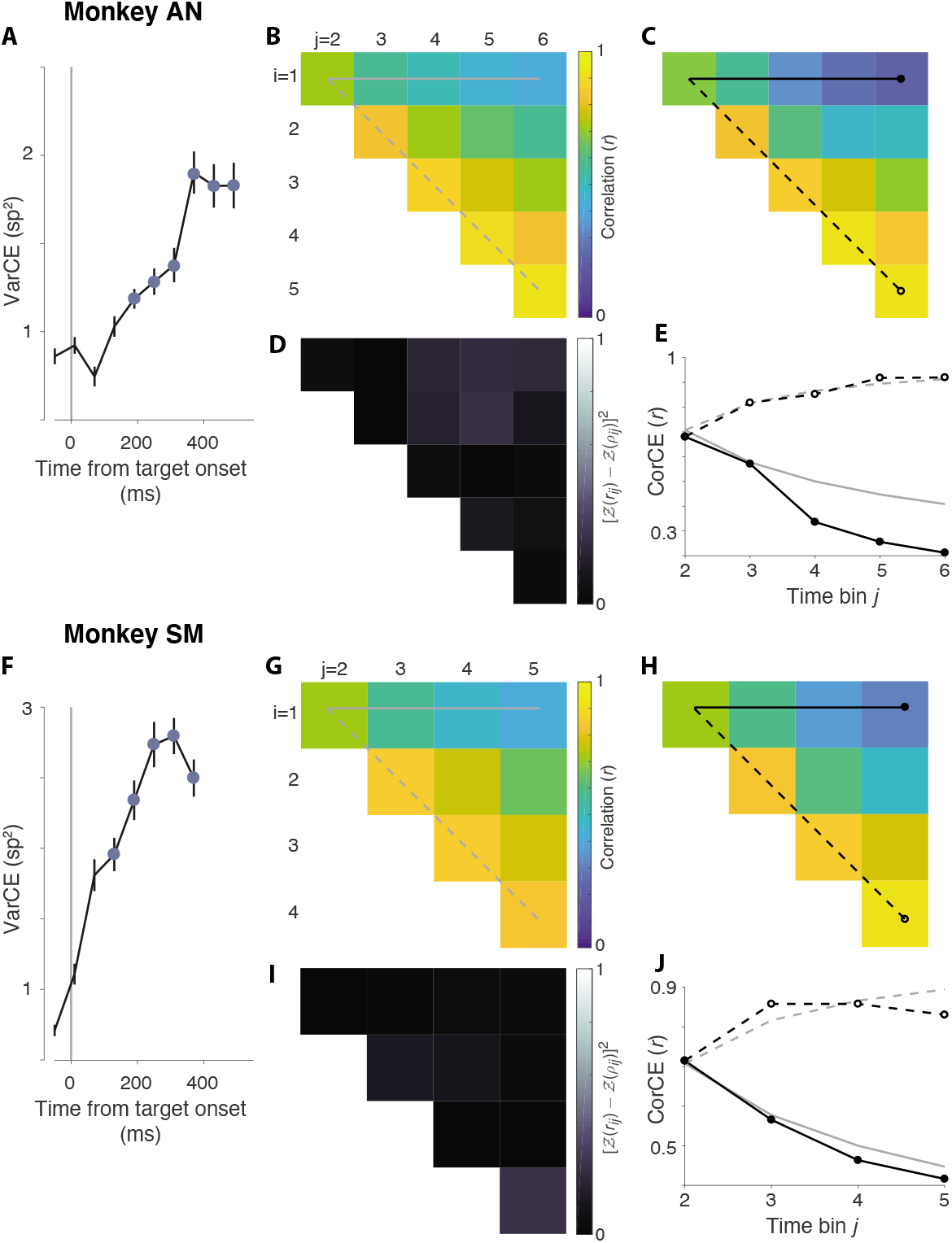
Variance and autocorrelation of decision related neural responses during action selection. The analyses depicted here evaluate predictions that the neural activity during action selection epoch on single trials includes a representation of accumulated noise. **A,F**, Variance of neural responses aligned to target onset. Filled symbols are estimates of the variance of the conditional expectations (VarCE) of the spike counts in 60 ms bins spanning the putative integration epoch. This is an estimate of the variance of the firing rate in the bin, across trials. **B,G**, Theoretical correlations between the cumulative sums of independent, identically distributed random numbers from the 1^st^ to *i*^th^ and from 1^st^ to *j*^th^ samples. The unique values of the correlation matrix are displayed as an upper triangular matrix. The horizontal solid line shows the correlation between the first sample and the cumulative sum to the *j*^th^ sample (lag = *j* – *i*). It shows decreasing correlation as a function of lag. The dashed line identifies the first juxtadiagonal set of correlations between pairs with the same lag = 1. It shows an increase in correlation as a function of time of the pairs of samples. The theoretical values for all correlations is given by 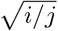. **C,H**, Correlations estimated from the neural response. These are the correlations between the conditional expectation of the spike counts (CorCE) in time bins *i* and *j*. This is an estimate of the correlation between the firing rates that gives rise to these counts. If the rates on single trials are determined by unbounded drift-diffusion, these correlations should match the values in panels B/G. The top row and first juxtadiagonal are identified as in B/G. **D,I**, Deviance of the estimated correlations from theoretical correlations (sum of squares measure). **E,J**, Comparison of theoretical and estimated correlations in the top row and first juxtadiagonal of the matrices in B/G & C/H. Gray traces show the theoretical values in B/G. Black lines connect the CorCE values in C/H. Line and symbol styles distinguish the top row (correlation as a function of lag) and first juxtadiagonal (correlations of neighboring bins as a function of time).

Importantly, analyses of the neural responses support the thesis that both monkeys form their decisions in the action selection epoch. Moreover, they do so through the accumulation of noisy samples of evidence to a threshold. The strategy is strikingly different from the previous studies [12,14] and it is all the more remarkable because it holds across two variants of the task. Below, we consider the reasons why our monkeys postpone decision making until the action selection. The fact that they do implies that their decisions are guided by evidence stored in short-term memory. This conclusion is at least equal in importance to the deferment of the decision process until action selection.

## Discussion

The study of perceptual decision-making in monkeys has provided insights into the process by which sequential samples of sensory evidence are accumulated over time [27, 32]. A peculiar observation in these studies is that the accumulation of evidence is instantiated by neurons associated with motor planning [21,31,33–35]. When monkeys are trained to indicate a decision between category A and B by action-1 and action-2, the neurons that represent the decision process are parsimonously characterized by their association with actions 1 & 2. This observation has led to the proposal that perceptual decision-making is embodied as a choice between potential actions [7, 8].

Yet monkeys can make perceptual decisions when they are unsure of the action that will be required of them to report their decision [12,14,15,36]. We set out to investigate how monkeys accumulate sensory evidence under these circumstances, and we did so using monkeys that had never learned an association between the decision and the action to report it. Instead they learned to associate leftward and rightward motion with the colors yellow and blue. No left-choice or right-choice targets were visible during motion viewing, but afterward, a yellow and a blue target were presented at unpredictable locations in the visual field and the monkey was required to choose one or the other color. We therefore predicted that they would not represent evidence in the form of an oculomotor plan but rather as a plan to invoke the appropriate rule: choose blue or choose yellow. Instead we found that the monkeys formed their decisions after the color-choice targets appeared—that is, during the period of action selection—after the source of sensory evidence had been extinguished.

Both monkeys based their decisions on samples of evidence that must have been retrieved from short-term memory. Monkey-AN developed this strategy spontaneously; monkey-SM did not, but appears to have adopted this strategy once we imposed a second waiting period during the action selection epoch. The striking change was evident in the longer time span of stimulus information used to inform decisions (Fig. 5B) to achieve a level of proficiency comparable to monkey-AN and many others we have trained on direction discrimination tasks. The go-RT from monkey-AN exhibited one peculiar feature. The difference in nondecision times for blue and yellow choices was nearly as long as the entire range of go-RT for either choice. The pattern suggests that monkey-AN makes a decision about blue and, failing to achieve sufficient support, switches to evaluating the evidence for yellow. This would seem absurd if stated as evidence for rightward and leftward motion, because evidence for rightward is evidence against leftward, and vice versa. However the sources of evidence bearing on the value of blue and yellow items do not typically have this antithetical relationship. We thus interpret the temporal offset in the blue and yellow go-RTs as a sign that the monkey makes two decisions in series (e.g., [20]) but is willing to terminate with a blue choice if there is sufficient evidence. This model is evaluated in Appendix S1. This interpretation is based on only one monkey so must be regarded as provisional.

For both monkeys, however, neural recordings from area LIP provided further confirmation that a sampling process transpired during the action selection epoch. On trials when one of the color-choice targets appeared in the neural response field, it produced a visual response plus a signal reflecting the direction and strength of the previously presented motion. The time course of the evolution was characteristic of an integration process—more specifically, the integration of noisy evidence acquired from the stimulus. While the memory requirements for the protracted integration of evidence may seem daunting, in Appendix S1 we show that it is not necessary to ‘replay’ the entire sequence of evidence samples during the action selection epoch; instead, storing a few samples of evidence is sufficient to achieve high levels of accuracy in the task (see also [20]).

Previous studies of perceptual decisions, dissociated from action, have not implicated a role for memory, but we suspect it played a role. An effective strategy to dissociate a decision from a plan of action exploits the delayed match-to-sample design [1], wherein a subject evaluates a sample stimulus and then, after a short delay, is presented with a second stimulus, which is compared to the first and classified as the same or different. It is assumed that the subject forms a categorical decision about the identity or category membership of the sample before the test stimulus is presented and therefore before an action associated with match and non-match can be planned. Using this approach, it has been shown that monkeys can report if the test and sample belong to the same category [1,2,4,37] or share similar properties like magnitude [38], numerosity [39] or speed/direction [40]. These studies focus mainly on neural activity in association cortex during sample and delay period. This activity often varies systematically with the relevant properties of the sample stimulus and is thus interpreted as a decision about the identity or category membership of the stimulus, independent of any planned action.

Our results suggest an alternative interpretation. Instead of a decision about category, the information about the stimulus might be encoded in short term memory to support a comparison with the test stimulus, critically, to establish the behavioral response. This decision may require multiple samples, but not if the match/nonmatch comparison is easy or if the sample (and test) stimuli do not supply multiple samples. A mechanism like this has been observed in the setting of a comparison of two vibrotactile flutter-vibration frequencies [41]. A related alternative is that the sample stimulus is processed as an instruction to brain circuits that organize the response to the test stimulus. The instruction might establish a criterion to classify the test or it might establish the appropriate sensory-response mapping. For example, test stimuli A and B might be associated with responses, match and non-match, respectively, if the sample is A, or irrespectively, if the sample is B. Such a mechanism has been documented in a simple olfactory delayed match to sample task in mice [42]. It does not require making a decision about the sample; it requires enacting a memory, cued by the sample, of the appropriate sensory-response mapping between test odor and behavioral response.

Two earlier studies of abstract perceptual decision-making used tasks similar to ours, but reached the opposite conclusion. The task in the Gold and Shadlen study [12] was nearly identical to ours but their monkeys failed to exhibit any signs of deliberation in the action selection epoch. The saccadic latencies were ~200 ms from color-target appearance, suggesting the monkeys had formed their decision about the color rule before the targets appeared. The only salient difference with the present study is that their monkeys had been trained previously to associate motion with eye movements to targets. We suspect that having learned to accumulate evidence for motion as an evolving plan to make a saccade, they were able to form a decision in another intentional way—for color rule instead of target location. A similar explanation applies to the study by Bennur and Gold [14]. Their monkeys made decisions in the presence of saccadic choice targets. In the version of their task that resembles ours (their Version 3) the monkeys were required to associate up and down motion with up and down targets or with down and up targets, depending on a colored cue presented after the motion had been shown. Decision-related changes in LIP activity were apparent during motion viewing and before the action-selection epoch. As in [12], the monkeys had been trained on a direct association between direction and an action and thus required only a slight elaboration: to switch the stimulus-response associations in accordance with the color cue.

Our result was anticipated by Wang and colleagues [15], who used a spatial integration task with separate evaluation and action selection epochs. The task structure used for one monkey resembles our *go*-task. It imposes a delay between the extinction of a static discriminandum and the presentation of the choice options. Similar to our monkey-AN, their monkey-T exhibited go-RTs that depended on the strength of the evidence experienced beforehand. They also report that the rate of rise of neuronal responses in Area PMd during action selection was dependent on stimulus strength. The Wang study also supports the hypothesis that sampling of evidence from memory may be necessary to form a perceptual decision when the evidence is provided before it is possible to accommodate it in an intentional context.

The near limitless capacity for abstraction in humans gives an impression of disembodied ideation. Humans can evaluate propositions about the world—what things are and what categories they belong to—without using them as objects of possible action. An alternative formulation, rooted in ecological perception [43], suggests that knowledge of the environment is in the service of what we might do, in the form of considerations and intentions [44, 45]. One activity humans pursue is reporting to other humans. The conversion of a provisional report to an action, like “look at the blue spot if the motion is rightward” permits humans to form a decision before the action is specified. The same logical structure applies to what monkeys—previously trained to associate rightward/leftward motion with an eye movement to the right/left—can achieve in abstract decision tasks like ours. If the human is not informed about the axis of discrimination until after the motion has been viewed, then like the monkey, humans too must rely on memory [46]. Further, studies of iconic short term memory demonstrate that such memory can be formed strategically in order to anticipate knowledge of the operations that may be required [16,47]. Thus both monkeys in our task must have learned to store the appropriate motion information in short-term memory buffers to enable action selection based on the colors of the choice targets.

The difficulty that our abstract decision task poses for naive monkeys might raise concerns about the relevance of our finding to human cognitive function. But consider. The simpler (non-abstract) version of the motion task invites an association between a source of evidence, derived from one part of the visual field, that bears on the relative value of options, instantiated by targets elsewhere in the visual field. It is representative of the type of foraging decisions that monkeys make naturally, but it is not in the repertoire of the animal’s experiences. They must learn that the relevant evidence bearing on action-selection is conveyed by a population of direction selective neurons with receptive fields that align to the patch of dynamic random dots. The same is true of a simpler blue-yellow decision task in which the color of an object near the point of fixation, say, determines whether a reward is associated with a blue or yellow choice target shown elsewhere in the visual field. The abstract decision in our task requires the animal to combine these types of decisions, either by (*i*) building a hierarchical decision in which the outcome of the motion decision substitutes for the colored object, to instruct the blue-yellow choice, or (*ii*) storing evidence from motion to resolve the subsequent color choice.

The hierarchical strategy is the one humans appear to exercise, as the effect of the strength of evidence on go-RTs is minimal in human subjects [48,49]. It is the way we would instruct another human to perform the task: “Decide whether the cloud of random dots is moving to the right or left; if right choose blue, and if left, choose yellow.” This is also the way monkeys previously trained on the non-abstract motion task solve the abstract task [12]. Yet both monkeys in the present study used the second strategy. Neither showed any sign of integrating evidence toward a decision during motion viewing. We speculate that this is because they never had the experience of planning an action associated with a decision about motion. They were rewarded only for actions associated with color, and they could only discover a source of evidence associated with the color selection in short-term memory. It is an open question whether they improved their performance by learning to store more samples or by sampling more from passive storage that occurs naturally during perception, or both.

At first glance, the hierarchical strategy might appear to be the more sophisticated of the two. It is more complex, and the nested structure seems like a building block for language. Indeed, humans probably adopt this strategy because it is implied in the verbal instruction to perform the task. However the second strategy also connects to a sophisticated element of cognition: the capacity to use recent, but temporally non-adjacent, information to guide a decision. This is critical for learning causal relations, and it too plays a role in language. We make strategic use of short-term memory to store semantic content (analogous to samples of evidence) which we incorporate in locution later—analogous to action selection—in accordance with syntactic demands. The process is embarrassingly vivid when we lose the train of our thought. Such embarrassment is mitigated by the strategic use of short term memory, adhering to the old adage, “Put your mind in gear before you put your mouth in motion” (A. Shadlen, *personal communication*).

Clearly, expressions of perceptual decisions through eye movements and expressions of ideas through language invite more contrast than comparison, but the structural similarity may prove useful for neurobiology. Thus it is that the monkey’s crude approximation to abstract decision-making elucidates a critical building block of our own ideation.

## Materials and Methods

All procedures were in accordance with the Public Health Service Policy on Humane Care and Use of Laboratory Animals, and approved by Columbia University’s Institutional Animal Care and Use Committee.

### Behavioral Task and Electrophysiology

Two adult macaque monkeys (one female, AN; one male, SM) performed a behavioral task in which they decided whether the net direction of a stochastic random-dot motion (RDM) stimulus was to the left or right. The animals initiated trials by fixating on a point (fixation point; FP) presented on an otherwise black screen. The RDM stimulus was then presented within a circular aperture (radius 2.5° or 3°) centered on the FP. The first three frames of the stimulus consist of white dots randomly plotted at a density of 16.7 dots·deg^-2^·s^-1^. From the fourth frame, each dot from three frames before is replotted—either displaced to the right or left, or at a random location. The probability with which a dot is displaced to the right or left determines the stimulus strength (coherence; *C*) and on each trial, *C* was randomly chosen from the set {0, ±0.04, ±0.08, ±0.16, ±0.32, ±0.64}, the positive sign indicating rightward motion. The motion strengths and the two directions were randomly interleaved. The stimulus was presented for a variable duration drawn from a truncated exponential distribution (range 350–800 ms, mean 500 ms). Two targets, one blue and one yellow, were presented after a short delay (333 ms, monkey-AN; 200 ms, monkey-SM) at eccentric locations that varied across trials. The monkeys had to report the perceived direction of motion by choosing the target of the associated color (blue for rightward and yellow for leftward, monkey-AN; vice-versa for monkey-SM). In the *go*-task (Fig. 1, top), the FP was extinguished simultaneously with the onset of the colored targets. In the *wait*-task (Fig. 1, *bottom*), the FP stayed on for a variable duration (drawn from an inverted truncated exponential distribution, range 400–1200 ms, mean 900 ms).

We recorded spikes from 60 well-isolated single units (29 monkey-AN; 31 monkey-SM) in area LIP_v_ [50]. The sample size for monkey-AN was limited by a serious illness, leading to euthanasia. Monkey-SM was just ready for recording when New York entered lockdown owing to the SARS-CoV2 pandemic. We justified as mission-critical the need to obtain a neural data set of power equivalent to the first monkey. The neural data were analyzed separately for each monkeys, and all of the central finding are statistically significant for each separately.

MRI was used to localize LIPv and to guide the placement of recording electrodes. We screened for neurons that exhibited spatially selective persistent activity using a memory-guided saccade task [25]. In the screening task, a target is flashed in the periphery while the monkey fixates on a central spot. The monkey has to remember the location of the target and execute a saccade to that location when instructed. The response field (RF) of each neuron was identified as the region of visual space that elicited the highest activity during the interval between the target flash and the eventual saccade.

During recording experiments, the locations for target presentation were chosen based on the location of the neuronal RF. For monkey-AN, six locations (including the RF) were chosen, equally spaced on an imaginary circle. On each trial, pairs of locations 2*π*/3 rad apart were pseudorandomly selected to display the targets. The RF location was oversampled to increase the concentration of trials from which we could analyze neural data. A similar approach was taken in monkey-SM except that the number of possible locations were restricted to four and the target pairs were situated *π*/2 rad apart. Each colored target appeared in the RF on 33% and 28% of the trials for monkey-AN and monkey-SM, respectively. Note that the monkeys were trained to generalize across a larger set of locations and these spatial restrictions on target locations were implemented during recording sessions.

### Analyses of behavioral data

Both monkeys were taught the association between the color of the target and the direction of motion using only the strongest motion strength (±64% coh). We then introduced the next easiest stimulus strength (±32% coh) and continued to add more coherences until we reached 0%. To assess the improvement of sensitivity across training sessions, we fit the choice-accuracy, *P*_correct_, as a function of motion strength, |*C*|, for each session with a Weibull function [51] of the following form:

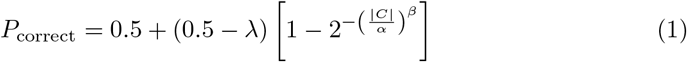

where λ is the lapse rate, *β* is the shape parameter, and *α* is the threshold if λ = 0. We interpolated from these fits the |*C*| that supports 75% accuracy and report that as the threshold (e.g., Supp. Figure 2-1).

The quantification of learning rate is from the introduction of the ±32% coh. The rates (e.g., Supp. Figure 2-1) are based on approximate number of sessions (and trials), because both monkeys experienced interruptions to training. For interruptions lasting more than a month, we excluded sessions after resumption until the monkey re-established thresholds similar to those prior to the interruption. This was also the case for monkey-SM when we switched from the *go*-task to the *wait*-task.

In Fig. 2A,B and Fig. 5C, we fit the choices of the monkeys with a logistic model of the following form:

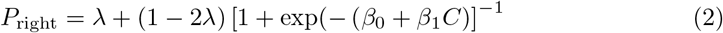

where λ, *β*_0_, *β*_1_ are fit parameters (Fig. 2A-B). This is also the analytic solution to symmetric diffusion (when λ = 0), and thus comparable to the fits of the models which are constrained to explain both choice and go-RT.

The go-reaction times (go-RT) of monkey-AN were fit with a bounded evidence accumulation model [52], modified to account for errors at the highest motion strength. In this model, the instantaneous evidence about motion at each time step is assumed to arise from a normal distribution with variance Δ*t* and mean *κ*(*C* + *C*_0_)Δ*t*, where *C* is the signed motion coherence, *C*_0_ is bias (expressed in units of signed coherence), and *κ* is a scaling parameter. The samples of instantaneous evidence are assumed to be independent and accumulated over time until the decision terminates, which occurs when the accumulated evidence reaches one of the bounds ±*B* leading to the choice of one of the targets. The mean go-RT is the expectation of the time taken for the accumulated evidence to reach the bound plus a constant—the non-decision time *t_nd_* comprising all contributions to the go-RT that do not depend on motion strength/direction and bias (e.g., sensory and motor delays). To account for asymmetric go-RTs in some configurations, we used two different non-decision times (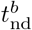 and 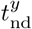) for blue and yellow target choices respectively.

In this framework, the mean go-RT for correct choices (i.e. choices consistent with the sign of the drift rate, *κ*[*C* + *C*_0_]) is described by

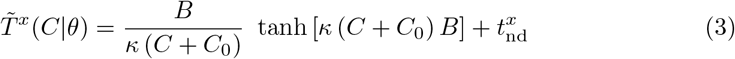

where *x* ∈ {*b, y*} and *θ* are the fitted parameters 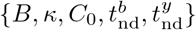. The proportion of blue choices is determined by three of these parameters:

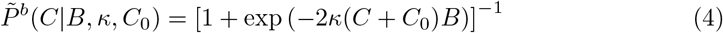

where 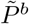 is the probability of the diffusion process terminating at the bound for blue choices. We first established an estimate of the bias from a logistic fit to the choices (Eq. 2), expressing the bias in units of coherence (*ζ* = *β*_0_/*β*_1_). Because the model explains the go-RT only when the choice is consistent with the sign of the drift rate [53], we used the mean go-RT for positive choices at *C* + *ζ* > 0 and negative choices for *C* + *ζ* < 0.

Informed by the patterns of error go-RTs observed at the highest coherence (Supp. Fig. 4-1B), we attribute the errors at the highest motion strength (lapse rate, λ) to a mistaken association between the sign of the terminating bound and its corresponding color-target (“direction-color confusion”). For weaker motion strengths the same confusion converts a fraction of correct terminations to erroneous color choices and the same fraction of incorrect terminations to correct color choices. We estimated λ from Eq. 2, thereby enabling conversion of 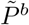 to the observed proportion of blue choices (*P^b^*):

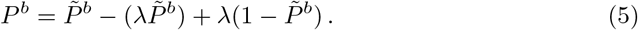

In our formulation, the trials with direction-color confusion inherit the *t*_nd_ of the motion decision (not the chosen color) and the mean observed go-RT would include contributions from the trials lost and gained from that process. The fraction of confusion trials for blue choices at coherence *C* is

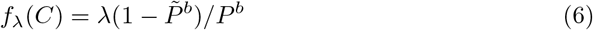

and the mean go-RT for blue choices observed to be correct would be

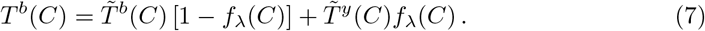

We used a maximum likelihood procedure to fit this model to the choice and mean go-RTs on the correct (relative to *ζ*) choices (Fig. 4). Table S1 shows the best-fitting model parameters. For each motion coherence, we calculate the average response time on correct trials (*RT^c^*(*C*)), its standard error (*s^c^*(*C*)), and the number of blue and yellow choices (*n^b^*(*C*) and *n^y^*(*C*) respectively). The parameters (Φ) are fit to maximize the function,

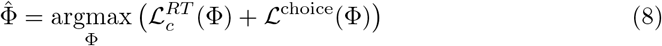

The first term of the right hand side of equation Eq. 8 is defined as:

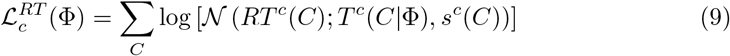

where *T^c^*(*C*|Φ) is the mean go-RT predicted by the model for motion strength *C* (correct trials only), and 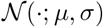 is the normal probability-density function with mean *μ* and standard deviation *σ*.

The second term of the right hand of equation Eq. 8 aims to maximize the probability of the observed choices given binomially-distributed errors,

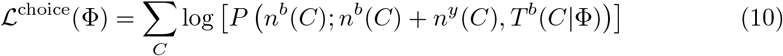

where *P*(*k*; *n,p*) is the binomial probability of observing *k* blue choices out of *n* trials, given that blue choices occur with probability *p*, which we take to be equal to the model’s predicted proportion of blue choices for parameters Φ.

We also fit an elaborated version of the bounded evidence accumulation model to include both correct and error trials. In this model, the decision bounds (*B*) collapse over time:

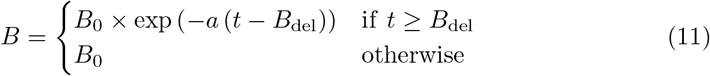

where *B*_0_ is the initial bound height, *a* is the rate of collapse and *B*_del_ is the delay to onset of collapse. The non-decision time was assumed constant and equal to *t*_nd_. Instead of using Eq. 3 and Eq. 4, 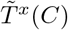 and 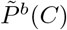 are obtained by numerical solution of Fokker-Planck equations ( [54,55]). Again, separate non-decision times were used for decisions terminating at each of the two bounds and errors at the highest coherence were modeled as ‘direction-color confusion’ using the approach described above.

We augmented these analyses with psychophysical reverse correlation, to provide an empirical estimate of the epoch in which the RDM stimulus affected the choice. The motion energy on individual trials (0% coherence only) was computed using spatiotemporal filters as described in [56]. The sign, right minus left or vice versa, was chosen such that positive indicates stimulus evidence in support of the monkey’s choice on that trial (Fig. 2C–D and Fig. 5B). To determine the actual duration of motion that had a significant influence on choices, we recalculated kernels using different lengths of the random dot movie shown in each trial. We report the length of time that the stimulus affects choice as the shortest movie-length that accounts for all the statistically significant bins obtained using the full-length movie.

### Analyses of neural data

For visualization of population average firing rates (Fig. 6), spike times from single trials, *s*_*i*=1...*n*_, were represented as delta functions *δ*(*s_i_* – *t*) and convolved with an 80 ms boxcar filter. For each neuron we grouped trials based on what was presented in its RF: blue target, yellow target or neither. We averaged across trials for each group and determined the maximum of the average responses across the three groups. The responses on all individual trials were divided by this maximum to obtain normalized firing rates. The population responses shown in Fig. 6 were then computed from these normalized responses using relevant subsets of trials. For the motion viewing epoch, trials were grouped based on motion direction (0° or 180°) and coherence (High: 64% & 32%; Medium: 16%; Low: 8% & 4%; and 0%). In the target onset and saccade epochs, the grouping was based on which target was shown in the neuron’s RF (blue or yellow), coherence (same coherence groups as in the motion viewing epoch) and the direction of motion (preferred vs. nonpreferred). For the majority of neurons, on trials in which a target appeared in the RF, a higher response was recorded when target appearance was preceded by the associated motion direction. For six neurons in monkey-SM, the non-associated direction elicited the higher response and was designated the preferred direction. To visualize the coherence dependent buildup of activity (insets of Fig. 6C–F), we detrended the population responses by subtracting the average responses to the 0% and ±4% coherence conditions. This detrending was done separately for trials with each colored target in the RF.

We pursued several analyses to characterize the neural responses during the epoch of action selection, after the onset of the color-choice targets. We defined the beginning of this epoch, *t*_∇_, as the first of three consecutive 40 ms time bins, beginning at least 50 ms after target onset, in which the average responses associated with correct choices at the strongest motion diverged (p<0.05, Wilcoxon rank sum test). For monkeys AN and SM *t*_∇_ = 170 and 100 ms, respectively. Our analyses focus on early decision formation, before many decisions would be expected to terminate on the more difficult conditions. For monkey-AN, we set the end of the epoch as *t*_∇_ + 300 ms or 200 ms before saccade initiation, whichever occurred first. There are no overt terminating events for monkey-SM. We therefore chose *t*_∇_ + 250 ms.

The effect of signed motion strength on build-up rate (Fig. 7) was established as follows. In the epoch defined above, we computed firing rates in 20 ms bins for each trial. For each neuron we grouped trials based on the target that appeared in the RF (blue or yellow). We removed the sensory component of the responses for each group by subtracting the average responses to the 0% and ±4% coherence conditions and computed the buildup rate for each coherence (the slope across bins). We excluded the ±64% coherence conditions from this analysis because there were too few time bins for monkey-AN, owing to fast go-RT, and an early plateau in monkey-SM, owing, we suspect, to fast decision terminations. We report the population mean and SE of the buildup rates and the fit to a linear model regressing these buildup rates against signed coherence in Fig. 7.

The analyses summarized in Fig. 8 compare the evolution of the variance and autocorrelation of the firing rate during the epoch of putative decision formation to the expected time course of these statistics under diffusion — if the spikes are associated with latent firing rates that represent the the sum of independent, identically distributed (*iid*) random numbers. The theory and algorithm are described in previous publications [29–31]. We used the spike counts in 60 ms bins in the epoch described above. This analysis focused on trials with the three weakest motion strengths (0%, ±4% and ±8% coh) to exploit the longer duration over which the decision process unfolds in these trials. The trials are initially grouped by neuron, the 5 unique signed coherences, and the target in the RF. We used the residuals of responses for each group to remove the contribution of motion strength and direction.

Consider, for the moment, trials from one neuron and one time bin. For each trial, *i*, we measure the raw spike count and compute the residual count by removing the mean count for all trials of the same combination of signed coherence and the color of the target in the RF, *j*,

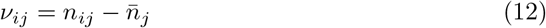

The total variance across trials, is

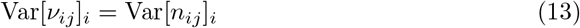

because variance is a central moment. We assume the noise component of evidence samples is the same for all the motion strengths. Therefore the variance across all combinations of signed coherence and the color of the target in the RF is Var[*v*], ∀ *j*. This is the total variance of the counts in the time bin under consideration. We are interested in the variance of the latent rate that gives rise to the spike counts on each trial. This is obtained by subtracting off the component of the variance attributed to the variable spike counts that would be observed even if the latent rate were fixed. For a Poisson point process this would be 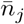, but we assume the point process is a generalized renewal [57] and is thus approximated by the *point process variance*,

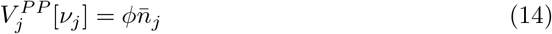

where the Fano factor, *ϕ*, is unknown. Note that the point process variance depends on the signed coherence. From the law of total variance, subtraction of this component from the total variance leaves the variance of the conditional expectation,

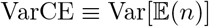. There is a bookkeeping step that respects the dependence of *V^PP^* on signed coherence and neuron (see previous citations), but using the residuals, we can obtain an estimate of the VarCE across all neurons at one time bin. Dividing by 0.06^2^ yields the variance of the latent rates (spikes^2^/s^2^) across trials (in the time bin under consideration), although it depends on the unknown *ϕ*. For unbounded diffusion the VarCE should increase linearly as a function of time, because it is a cumulative sum of *iid* random numbers.

Diffusion also specifies the autocorrelation, between the cumulative sum of the first *i* samples and the cumulative sum of the first *j* ≥ *i* values:

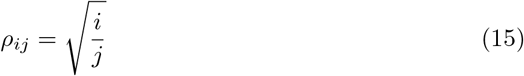

This implies a decay of correlation as function of lag, *j* – *i*, and an increase in correlation for fixed lag, as a function of time. We obtain the estimates, *r_ij_*, from data by forming the autocovariance matrix on residuals from all neurons. Note that the *Cov^PP^* = 0 for *i* ≠ *j*, because by construction, given the rate in time bin *j* the stochastic realization of spike count does not depend on the rate or realization of spike count in bin *i*. Therefore the covariance of the conditional expectation (CovCE) is the raw covariance for *i* ≠ *j*. Its diagonal (*i* = *j*) is the VarCE. This matrix is normalized in the usual way to produce a correlation matrix of conditional expectation (CorCE).

The CorCE depends on the VarCE which depends on *ϕ*, which is unknown. We chose the value that minimized the sum of squares,

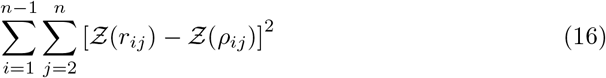

where 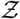 denotes standardization (Fisher-z transform).

The values of the variance plotted in Fig. 8A,F are VarCE, using the fitted *ϕ*. The standard errors are estimated from a bootstrap procedure [58] in which trials were sampled (with replacement) while maintaining their grouping (same neuron, dot direction, coherence and color of target in RF). We also performed the same analysis using neural responses in the epoch between 190 to 550 ms after RDM onset (Supp. Figure 6-1 & Supp. Figure 7-1).

## Acknowledgements

The research was supported by HHMI, NIH and BBRF. We thank Brian Madeira and Cornel Duhaney for technical support and animal care. We thank Danique Jeurissen, Gabriel Stine, Natalie Steinemann, Simon Kelly and Redmond O’Connell for comments on an earlier draft of the manuscript, and Arthur Shadlen (grandpa) for advice and/or reprimand. We are especially grateful to animal technicians, veterinary staff, and other essential workers at the Zuckerman Institute who made it possible to collect the data set from monkey-SM during the SARS-CoV-2 pandemic.

---

**Table S1.** Parameters of the best-fitting model shown in Supp. Fig. 4-1.

| Parameter | Description | Best-fit |
| --- | --- | --- |
| $\kappa$ | Signal-to-noise | 12.1 |
| $t_{\text{nd}}^b$ | Mean non-decision time for blue/rightward choices | 0.37 |
| $t_{\text{nd}}^y$ | Mean non-decision time for yellow/leftward choices | 0.56 |
| B | Bound height | 0.46 |
| C0 | Bias | -0.01 |
| $\lambda$ | Probability of direction-color confusion | 0.09 |

**Supp. Figure 2-1:**
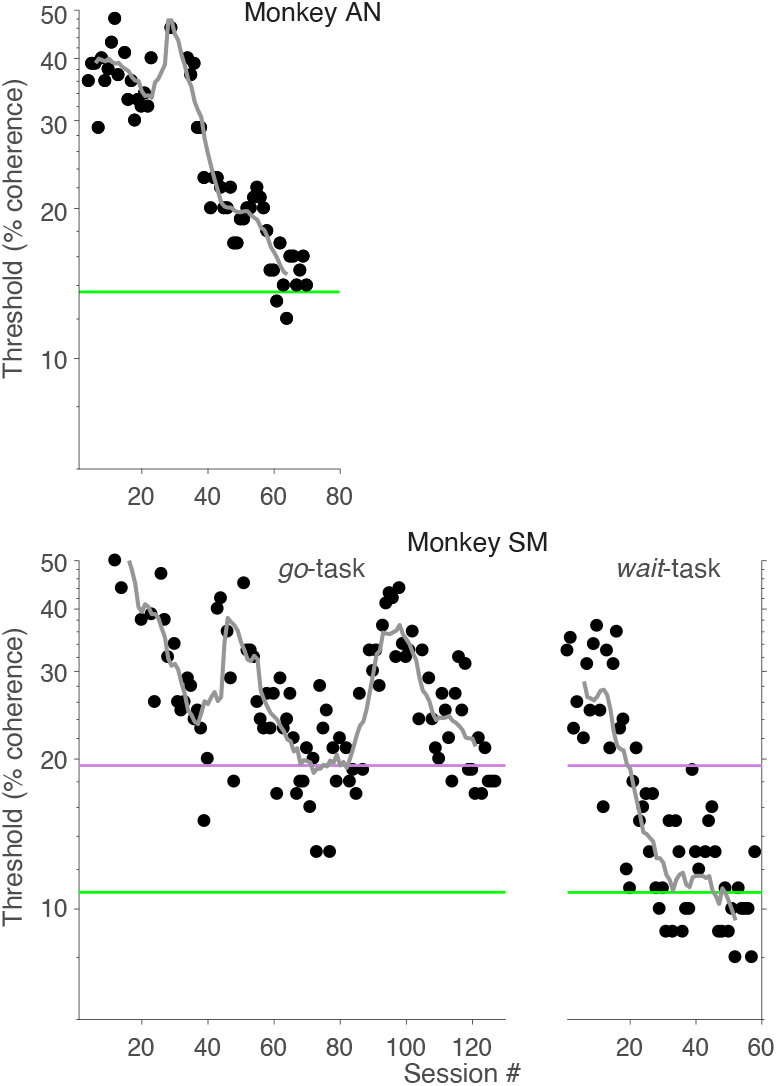
Psychophysical thresholds during training. The motion strength required to support 75% accuracy is plotted as a function of training day. The values are interpolated from a Weibull fit (Eq. 1) to proportion accuracy vs. |coherence|. Green lines indicate the thresholds from all neural recording sessions. The purple line in the bottom panel indicates the thresholds for monkey-SM from the last four sessions of training on the *go*-task (the sessions included in Fig. 5). Grey lines show the running geometric mean of thresholds from 11 consecutive sessions.

**Supp. Figure 4-1:**
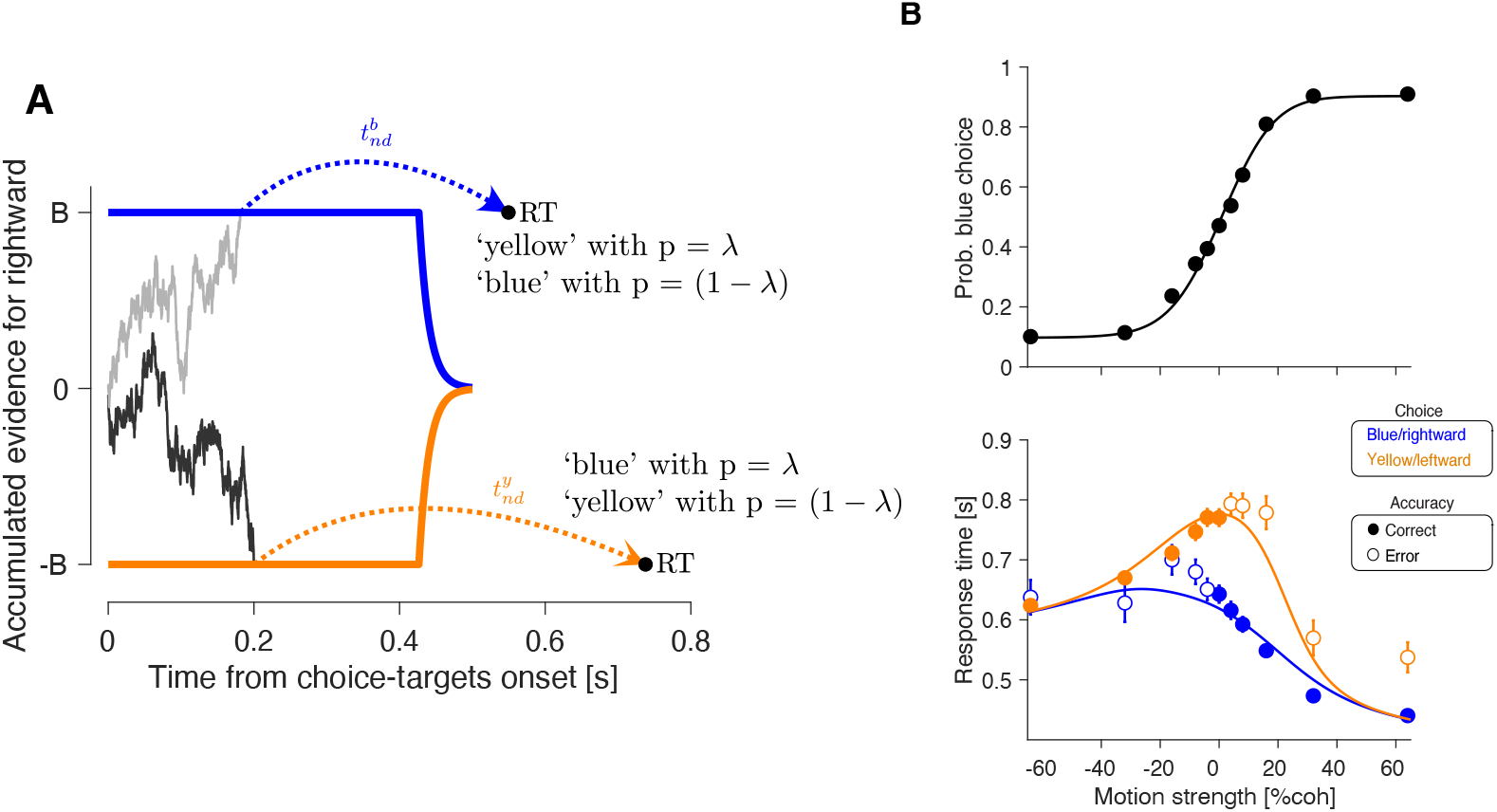
A decision model with collapsing bounds and direction-color confusion explains the go-RTs on errors. We fit a drift-diffusion model with collapsing decision bounds to the choice and go-RT data, including the error go-RTs. (A) Sketch of the drift-diffusion model with collapsing bounds. Two trials are shown, terminating at the upper and lower bounds (gray and black traces, respectively). A fraction of trials, λ, deploy the wrong association between motion direction and color. The nondecision time (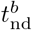 or 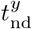) is associated with the sign of the terminating bound, so opposite that of the rendered choice on these trials. The sketch of the drift-diffusion model was generated using the parameters that best-fit the data (Table S1). The bounds collapse rapidly after ≈400ms, limiting the duration of the integration period. (B) Choice and go-RTs plotted as a function of signed motion coherence. Curves are fits of the drift-diffusion model with collapsing decision bounds and direction-color confusion, depicted in panel A. Error bars indicate s.e.m. across trials.

**Supp. Figure 5-1:**
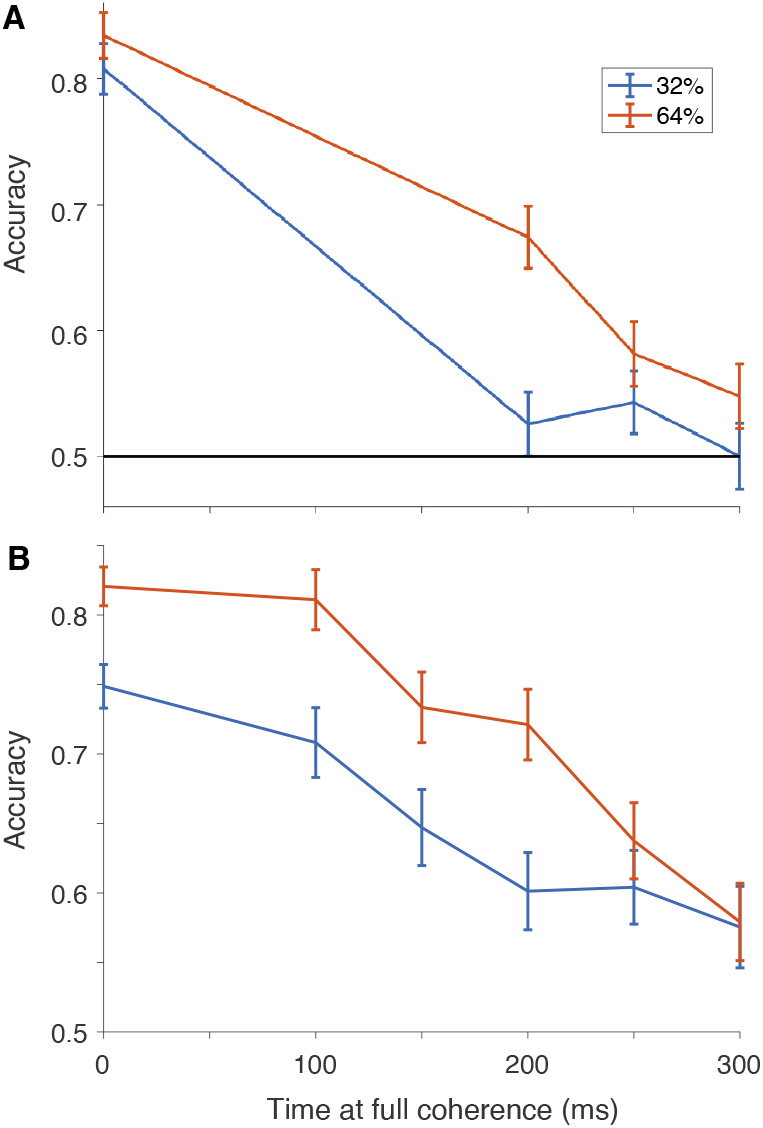
Further confirmation that monkey-SM did not accumulate evidence from the motion stimulus on the *go*-task. We introduced two variants of the *go*-task to evaluate the hypothesis that monkey-SM used only a brief sample of the random dot stimulus to inform its choice. **A**, Step version. Each trial started at 0% coherence and stepped to ±32% or ±64% at a variable time. Choice accuracy is plotted as a function of step time (4 sessions, N=4663 trials). **B**, Ramp version. The motion strength changed linearly as a function of time from 0% to ±32% or ±64% coherence. The rate of change was varied across trials. Choice accuracy is plotted as a function of the time when the coherence reached its maximum (3 sessions; N=3054 trials).

**Supp. Figure 6-1:**
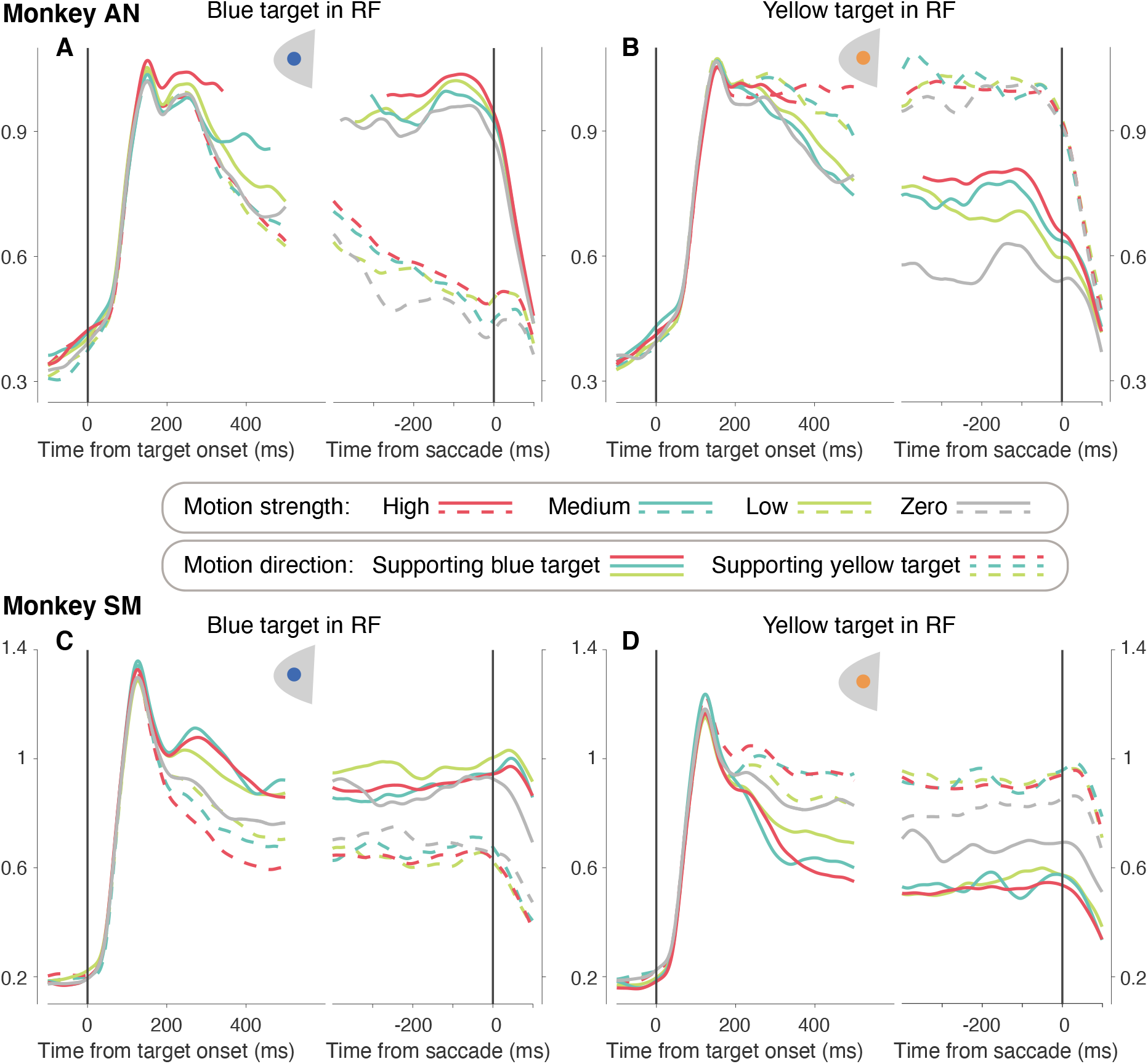
LIP activity during target selection. The graphs show average normalized responses as a function of time aligned to target onset (left half of each panel, same data as in Fig. 6) or time of saccade (right half of each panel). For the responses aligned to target onset, all trials, including errors, are included in the averages grouped by signed coherence; for the responses aligned to saccade, errors are excluded. Rest of the conventions are as in Fig. 6.

**Supp. Figure 8-1:**
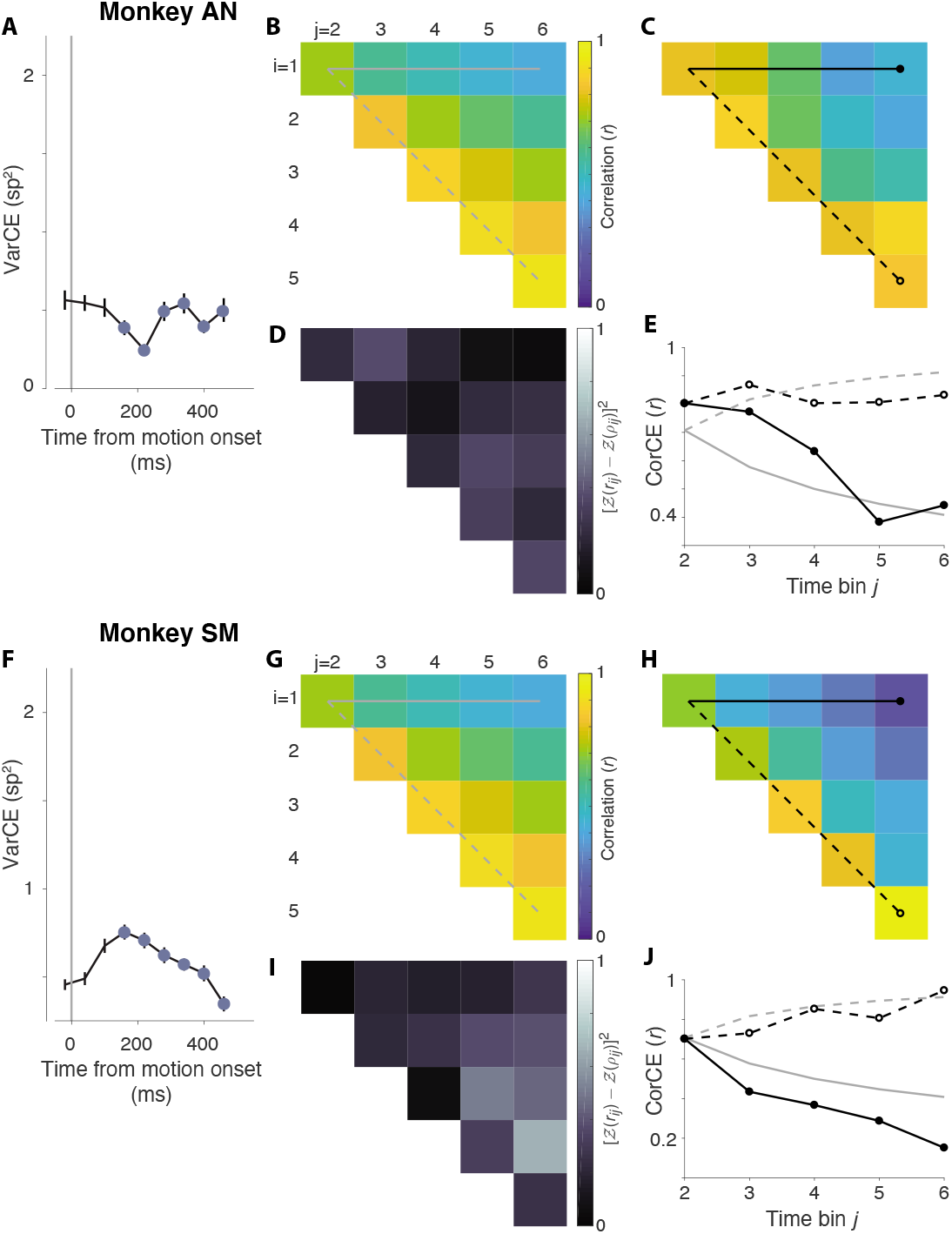
VarCE and CorCE during motion viewing. Same analysis as in Fig. 8, using data from the motion viewing epoch.

## Appendix S1: Additional analyses of go-RTs from monkey-AN

### 1 Overview

The pattern of go-RTs of monkey-AN provides additional insights into the decision-making process. In this appendix, we elaborate on two intriguing aspects.

1. The relatively high rate of errors at the strongest motion strength.
2. The asymmetry in the go-RTs to the two colored targets.

Further, we speculate on how motion information can inform a decision process that commences hundreds of milliseconds later.

### 2 Low sensitivity cannot account for errors on easy trials

We have hypothesized that a substantial fraction of errors—indeed the vast majority of errors at the highest motion strengths—are explained by a mistaken association of direction with color. An alternative is to attribute these errors to a noisier or less efficient decision process, perhaps because the evidence must be held in memory.

To distinguish between these alternatives, we compared the drift-diffusion model described in the main text to one in which the errors at the highest coherence are not explained by direction-color confusion but by low sensitivity. Here we refer to these two models as the two-*t*_nd_ model and the *Low Sensitivity* model, respectively. Both models are identical except except that in the Low *Sensitivity* model the probability of a direction-color confusion (λ) was set to zero.

Model comparison provides decisive support for mistaken associations of direction and color. Fig. A1 overlays the fits of both models. The dashed line shows the best fits for the *Low-Sensitivity* model (λ = 0). The difference in log-likelihood between the best-fits of both models was 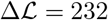 in favor of the model with non-zero λ. It thus appears that on the vast majority of trials the monkey forms a decision based on the accumulation of evidence from short-term memory and makes the opposite color choice on ~10% of trials.

### 3 Decision model with prioritization

The go-RTs of monkey-AN are strikingly asymmetric (Figure 4), such that the entire range of mean go-RTs is approximately the sum of the ranges for either choice. In the model accompanying the main text, we accommodated the difference in the two non-decision times, but raised the possibility that the asymmetric pattern might reflect a serial process.

**Fig A1.**
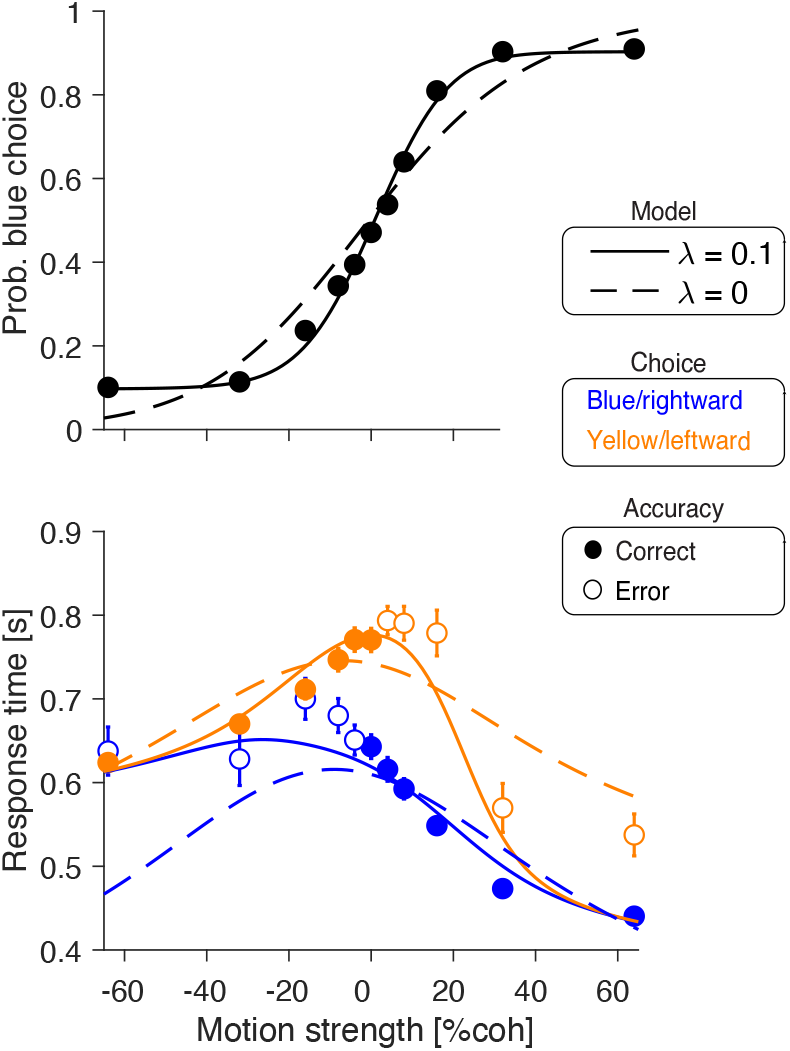
Comparison of models with and without direction-color confusion. We consider two explanations of the errors at the highest motion strengths. Both models use bounded drift diffusion with symmetric, collapsing bounds. The **two-t**_nd_ model (solid traces) was explained in the main text. On a fraction of trials, λ, the decision-maker deploys the wrong association between motion direction and color. The **Low-Sensitivity** model (broken traces) implements the hypothesis that lapses are a sign of low sensitivity to motion information. This is the diffusion model without direction-color confusion (i.e., λ = 0). The two-*t*_nd_ model is clearly superior to the *Low Sensitivity* model. Data points correspond to data from monkey-AN. Error bars are s.e.m. across trials. The best-fitting parameters are reported in Table A1.

Here we consider a model in which the evidence for blue and yellow is evaluated serially [1]. In any one trial, the decision maker prioritizes the evaluation of the blue or yellow choice. The evaluation consists of the accumulation of noisy momentary evidence until reaching an upper bound (at *B*) or until reaching a deadline (*t*_deadline_) (Fig. A2A). If the upper bound is reached, the decision maker commits to the prioritized alternative, say blue, as depicted in Fig. A2A. If the upper bound is not reached by *t*_deadline_, this is interpreted as a failure to achieve sufficient support for blue, and the decision maker commits to the non-evaluated choice, yellow. Fig. A2A illustrates the dynamics of the model with two hypothetical trials, one that terminates at the upper bound and one that does not. As above, the actual choice is flipped on a fraction, λ, of the times, owing to a mistaken association between motion direction and color.

Unlike most evidence accumulation models used to account for binary choices, our model does not include a lower decision-termination bound. The absence of a lower bound implements the intuition that at the level of sensory processing, the evidence against blue is not necessarily evidence for yellow, nor vice versa. This is different from the way motion processing is organized in the primate brain, where evidence against rightward is evidence for leftward.

The probability density function for terminating the decision at the bound has a closed-form solution given by an inverse-Gaussian distribution [2]:

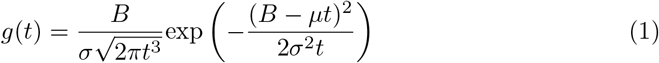

where *μ* and *σ* are the mean and standard deviation of the accumulated evidence after 1 second of boundless stimulus viewing. In our application, the mean is a function of the motion strength and bias, *μ* = *κ*(*C* + *C*_0_), and 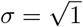, respectively. The bias *C*_0_ takes value 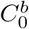 when evaluating the evidence for the blue choice, and 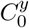 when evaluating the yellow choice. *C* is positive for rightward motion and negative for leftward motion, when evaluating the blue choice; conversely, it is negative for rightward and positive for leftward when evaluating the yellow choice. Therefore, when the evidence is strong toward the alternative being evaluated, the deterministic component of the drift-diffusion process is toward the bound.

**Table A1.** Best-fitting parameters for the two-*t*_nd_ and *Low-Sensitivity* models

| Parameter | Description | two-$t_{nd}$ model | Low-sensit. model |
| --- | --- | --- | --- |
| $\kappa$ | Signal-to-noise | 11.81 | 4.25 |
| $t_{nd}^b$ | Non-decision time for blue choices | 0.37 | 0.21 |
| $t_{nd}^y$ | Non-decision time for yellow choices | 0.54 | 0.43 |
| B0 | Bound height before collapse | 0.52 | 0.59 |
| a | Bound decay rate | 72.21 | 10 |
| $B_{del}$ | Time at which bound starts to collapse [s] | 0.43 | 1.38 |
| C0 | Bias | -0.01 | 0.04 |
| y0 | Starting point bias | 0.06 | -0.11 |
| $\lambda$ | Prob. of a direction-color confusion | 0.1 | N.A. |

Fig. A2B shows the fits of the model to the choice and go-RT data. The model provides a good fit to both the choice and go-RT functions. The model misses the go-RTs for error trials of strong rightward motion, for reasons that will become clear below.

To convey the intuition of why this simple model is capable of accounting for behavioral data, we dissect the model by showing its predictions after removing some of the model parameters.

Fig. A2A shows two hypothetical trials of the same motion strength, one that terminates at the upper bound and one that terminates at the deadline. In these hypothetical trials, the evidence for blue is evaluated first. Average response times are shown by the solid lines in Fig. A2C. Here we assumed that λ (the probability of direction-color confusion) is zero, and thus all left choices have the same go-RT, equal to the deadline (*t*_deadline_) plus a constant that represents the non-decision latencies (*t*_nd_). Blue choices, in contrast, depend on the time it takes the diffusing particle to reach the upper bound and thus the RT is modulated by motion strength. Likewise, when yellow is evaluated first, yellow choices are faster than blue choices (Fig. A2D).

The combination of trials that begin by evaluating the evidence for blue and those that begin by evaluating yellow leads to a pattern of go-RTs that is closer to what we observe in the data. The probability of starting evaluating the blue option, 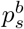, is a free model parameter that is fit to the data. The full model is obtained by allowing λ to be non-zero; λ is the probability of a mistaken association between the sign of the terminating bound and its corresponding color-target (“direction-color confusion”). This parameter explains the lapses observed at high motion strength and helps approximate the pattern of go-RTs observed in the data (Fig. A2B). The approximation, however, is not perfect, as there is a clear miss in the go-RTs for errors made at strong rightward motion. The model predicts that errors at high coherences have the same decision time as the correct choices at the same signed coherence. While this is true for strong leftward motion, it is not for strong rightward motion. We do not have a good explanation for this discrepancy.

**Fig A2.**
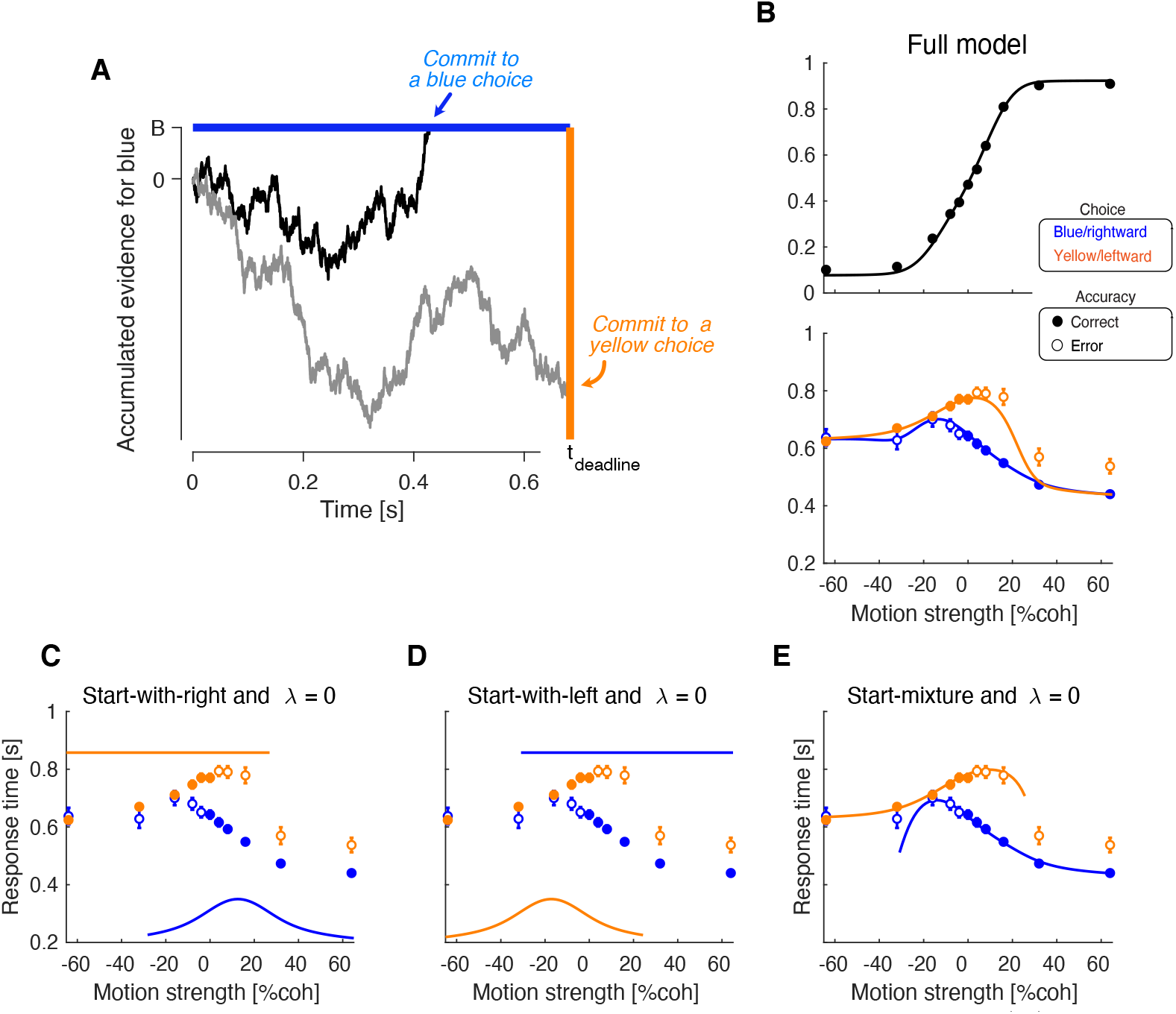
Simulations and fits of the prioritized integration model with deadline. (A) Sketch of the model. Two trials are shown. The decision process stops when the diffusing particle reaches an upper bound or a deadline. Because in these example trials the decision maker prioritizes the evaluation of the blue choice, the upper bound represents the commitment to a blue choice. If the upper bound is not reached by the deadline, the decision maker commits to a yellow choice. The model also allows for the possibility of color-confusion errors after the commitment to either choice. (B) Best-fits of the model. (C) Data is overlaid with the model predictions generated using the best-fitting parameters after setting λ to zero and 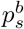 (the probability of starting with the blue choice) to 1. (E) Same as panel C, but for 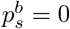 (i.e., yellow evaluated first). (E) Model predictions using the best-fitting model but setting λ to zero. In panels B–E, the go-RT predictions are not shown if the model chose an option less than 2 % of the times.

The model we present in the main text to explain the go-RT pattern expressed by monkey-AN assumes that blue and yellow choices have different non-decision times. Here we refer to it as the ‘two-*t*_nd_’ model. Although we consider this model to be a proxy for a model with prioritization such as the one we presented above, we performed a model comparison to ask whether the prioritization model better explains the data than the two-t_nd_ model. To achieve this, we compare the model with prioritization to the model with separate non-decision times for blue and yellow choices and a collapsing decision bound, which we presented in the main text.

Because the two models have the same number of parameters (see Table A2 and Table A1), we can compare their goodness-of-fit by directly comparing their likelihood. This comparison strongly favored the model with prioritization over the model with separate non-decision times 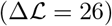.

**Table A2.** Best-fitting parameters for the model with prioritization

| Parameter | Description | Best fit |
| --- | --- | --- |
| $\kappa$ | signal-to-noise | 14.95 |
| $B$ | Bound height | 0.31 |
| $t_{\text{deadline}}$ | Deadline | 0.68 |
| $t_{\text{nd}}$ | Non-decision time | 0.17 |
| $C_0^b$ | Bias when evaluating the blue choice | -0.13 |
| $C_0^y$ | Bias when evaluating the yellow choice | -0.17 |
| $\lambda$ | Prob. of a direction-color confusion | 0.077 |
| $p_s^b$ | Prob. of starting evaluating the evidence for blue | 0.65 |

**Fig A3.**
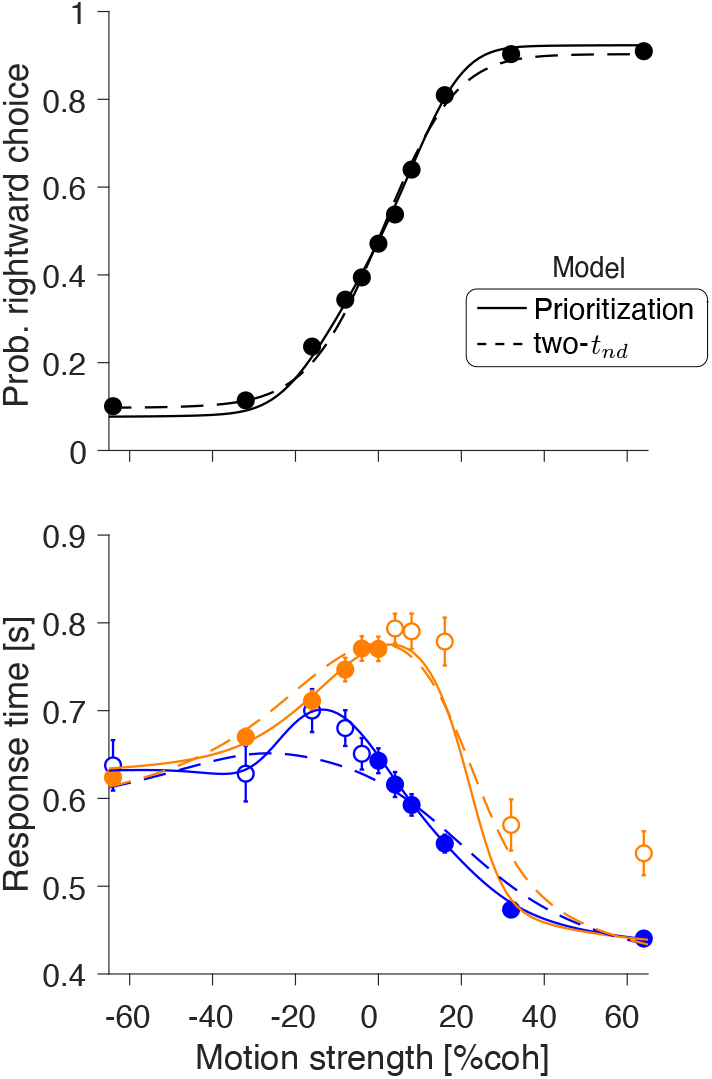
Data and fits of the prioritization and two-*t*_nd_ models. Solid and dashed lines depict the best-fit of the prioritization and two-*t*_nd_ models. While both models accurately fit the choice function, the go-RTs are better explained by the model with prioritization.

Fig. A3 overlays the best-fits of the two models. Both provide excellent fits to the proportion of blue choices as a function of motion strength, but the prioritizing model explains the go-RT pattern better, for both correct and incorrect options. Therefore, the difference in go-RT between the blue and yellow options displayed by monkey-AN may be the result of prioritizing the evaluation of the evidence for one of the options (‘blue’ in this case). Implicit in this explanation is the limited and serial access to information in memory, which is consistent with extensive psychophysical literature in humans [3,4], and more recent studies in non-human primates [5].

### 4 How can motion information be carried forward in time?

Integrating a stream of motion information hundreds of milliseconds after it disappeared from the display may seem to require a complex mechanism. One such mechanism is to store the full stream of information present in the motion stimulus, and replay it when the choice alternatives are revealed. It is not clear, however, whether the primate brain is able to perform such a feat.

**Fig A4.**
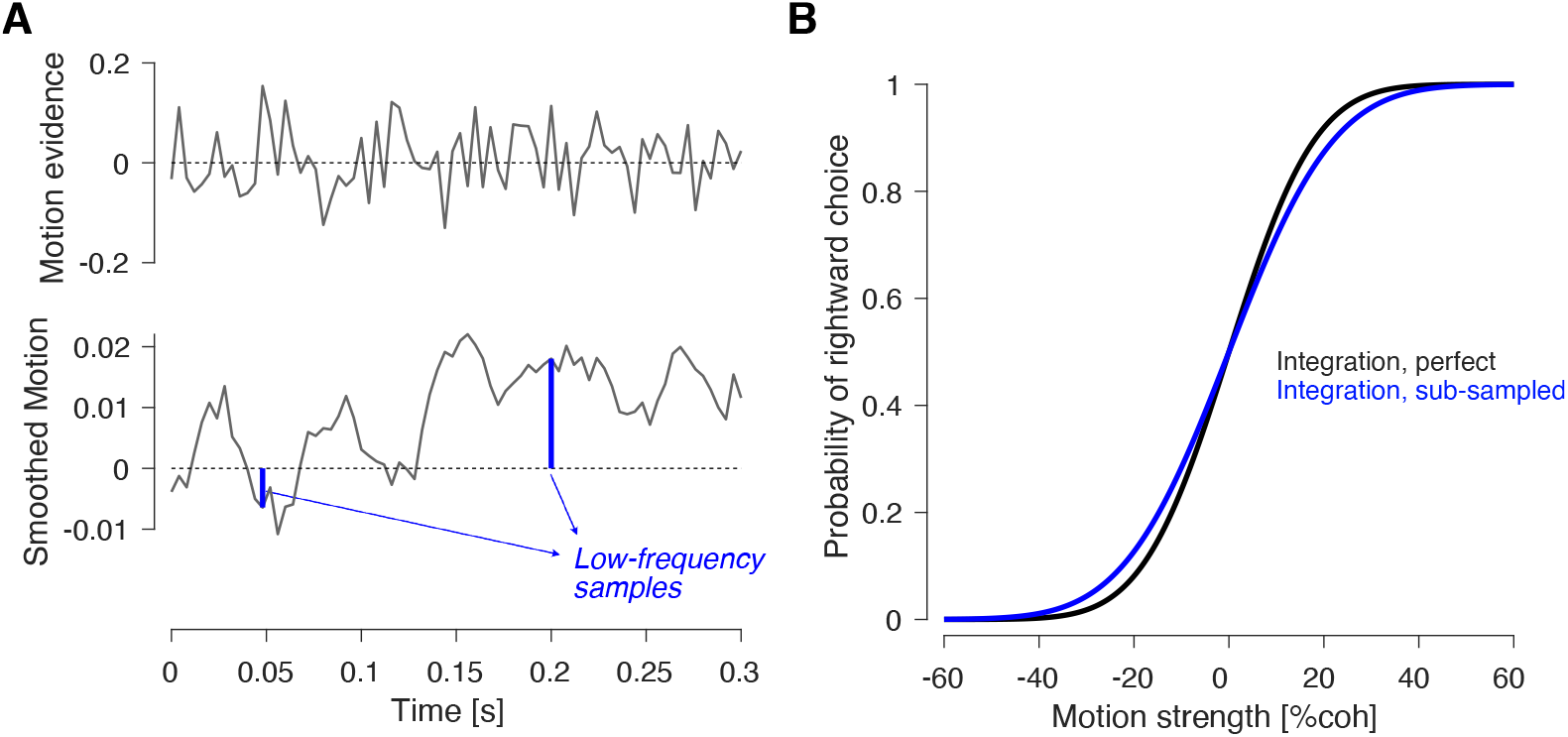
Comparison of decisions based on perfect and coarse integration of motion information. (A) The top panel shows hypothetical samples of the motion information present in the stimulus, assumed to be Gaussian with a mean of *κC*Δ*t* and a variance of 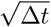. Here *C* = 0.2, Δ*t* = 4 ms, and *κ* is obtained from the fits of the two-*t*_nd_ model (Table A1). For the lower panel we applied a boxcar filter with a width of 100 ms to the trace above. We show two samples, separated by 150 ms. (B) Predicted proportion of rightward (blue) choices under perfect integration (black) and sub-sampled integration (blue).

Here we consider a much simpler alternative. It involves taking samples of motion information at a low frequency—that is, low relative to the sampling rate required to extract motion information from the video display—and using these samples as the basis for the decision. The process is illustrated in Fig. A4A. The momentary evidence present in the stimulus fluctuates rapidly, which we have represented as samples from a Gaussian distribution with a mean that depends on the trial’s motion strength (Fig. A4A, top). The motion computation in the visual cortex introduces blurring (autocorrelation) thereby supporting sub-sampling of motion information by downstream processes (Fig. A4A, bottom). The subsampling remains sufficient to capture the stochastic fluctuation in motion information that occur during each RDM movie. Fig. A4B shows the expected psychometric functions for a decision based on 300 ms of motion information, that is based on perfect integration of the motion stimulus (black trace), or the integration of motion information that is sub-sampled; in the figure, motion samples are taken every 150 ms (blue trace).

